# Persistent circulating autoreactive PD1⁺TIGIT⁺ peripheral helper T cells reflect synovial lymphoid activity and poor response to conventional disease-modifying anti-rheumatic drugs in early rheumatoid arthritis

**DOI:** 10.64898/2026.07.28.740434

**Authors:** Jia Yi Hee, Hendrik J Nel, Oxana Radetskaya, Gabrielle Antonio-Carreon, Carl Coyle, Wittaya Suwakulsiri, Megan Soon, Pascale Wehr, Mai T Tran, Garrett Dunlap, Deepak Rao, Annabelle Small, Soon Wei Wong, Katrina Chakradeo, Michelle Roch, Tom Lynch, Lyn M March, Andrew P Cope, Jamie Rossjohn, Mihir D Wechalekar, Yann Abraham, Ranjeny Thomas

## Abstract

**Objective:** In the first year after onset of the autoimmune disease RA (RA), 40-60% do not achieve remission on conventional synthetic disease-modifying anti-rheumatic drugs (csDMARDs). To understand how autoreactive T cells may contribute to unstable or non-remission, we studied CD4+ T cells, including those recognising citrullinated (Cit) vimentin in participants with RA.

**Methods:** Two cohorts of drug-naïve new-onset participants with RA were treated with csDMARDs. Disease activity score (DAS28-CRP) and peripheral blood (PB) mononuclear cells were collected longitudinally. In HLA-DR-shared epitope+ cohort 1 (n=21), T cells were assessed with a 17-marker spectral flow panel, incorporating HLA-DRB1*04:01/01:01- VimentinCit64_59-71_ or HLA-DRB1*04:04-VimentinCit71_66-78_ tetramers. Changes in T cell subsets over time were assessed in remitting and non-remitting participants using a generalized linear mixed model with a negative binomial distribution. In cohort 2 (n=26), the transcriptome of disaggregated synovial tissue (ST) and PB CD4+ T cells were analysed at baseline, and ST biopsy spatial proteomics at baseline and 6 months.

**Results:** CD4⁺CXCR5^-^PD1^+^ peripheral helper T cells (Tph), including Cit-vimentin-reactive Tph, CD4⁺CCR7^+^CXCR5^-^PD1^+^ stem-like Tph and TIGIT+PD1+ Tph were increased with moderate/high DAS28-CRP at any time point. Remission outcome was associated with low CD4⁺ follicular helper T cell (Tfh) and Cit-vimentin-reactive Tfh. In non-remitting participants, Tph/fh infiltrated germinal-centre-like ST aggregates. This decreased in remission. Circulating TIGIT^+^ Tph genes reflected B lymphoid activation and lymph node egress, while in ST they reflected local differentiation.

**Conclusion:** Persistently high circulating TIGIT^+^ Tph and Tfh, including Cit-vimentin specificities, reflecting antigen-presenting B-cell interactions, are associated with reduced response to csDMARDs in recent-onset RA.

**Key messages:** *What is already known on this topic:*

- Tph and Tfh cells interacting with B cells are implicated in active ACPA+ RA
- Citrullinated (Cit)-vimentin is an important neutrophil extracellular trap-derived target of ACPA

*What this study adds:*

- High circulating TIGIT+ Tph, Cit-vimentin-autoreactive Tph and Tfh associate with failure to reach remission on conventional synthetic DMARDs within the first year in early HLA-DR shared-epitope+ RA
- Tph/Tfh and adjacent regulatory T cells surround follicular B-cells in lymphoid aggregates in synovial tissue in non-remission
- Circulating TIGIT+ Tph bear a transcriptional signature of lymphoid tissue expansion and lymph node egress as compared to functional differentiation and antigen experience in synovial tissue

*How this study might affect research, practice or policy:*

- When a remission target is not achieved in HLA-DR shared-epitope+ RA patients during the first year of treatment, high circulating TIGIT+ Tph implicate autoreactive T-B-lymphoid expansion in synovial tissue.

## Introduction

Rheumatoid arthritis (RA) is a systemic autoimmune disease characterized by immune-mediated destruction of the synovial joints. RA is associated with the human leukocyte antigen (HLA)-DRB1, which encodes a component of the major histocompatibility complex (MHC) class II molecule responsible for presenting self- and non- self-antigens to CD4⁺ T cells (1, 2). Seropositive RA is particularly associated with HLA-DRB1 alleles encoding the shared epitope (SE), commonly including HLA-DRB1*04:01, HLA-DRB1*04:04, and HLA- DRB1*01:01 in Caucasians (3, 4). Remission in early RA is associated with favourable long-term outcomes, particularly when sustained over time, including reduced joint damage, and preserved physical function (5). Conversely, persistent disease activity or unstable remission is associated with progressive structural damage and irreversible disability (5). Despite treat-to-target strategies, clinicians lack reliable early biomarkers in the blood that identify patients requiring treatment escalation or modification to achieve and maintain remission, as well as evidence of which pathological processes they need to target in the synovial tissue (ST).

Citrullinated (Cit)-vimentin is an important neutrophil extracellular trap (NET)-derived target of anti-Cit peptide autoantibodies (ACPA) (1). Certain cell states of various CD4+ T cells recognising citrullinated self-antigenic peptides presented by HLA-DR-SE molecules may reflect and contribute to immune tolerance, RA disease development, activity and treatment response (2–6). However, the scope of markers has been limited and a broader understanding, particularly of CD4⁺ peripheral (Tph) and follicular helper T cells (Tfh) that interact with autoreactive B cells to induce maturation for antigen presentation and antibody production, is needed (7). Furthermore, although various Tph and Tfh cell states interacting with B cells are implicated in ACPA+ and ACPA- RA (8–11), it is not known how autoreactive CD4+ T cells contribute to the relationship between Tph in circulation (peripheral blood, PB) and synovial tissue (ST) in clinically active or remission states.

To understand the potential contribution of CD4+ T cells and CD4+ T cells recognising common Cit-vimentin epitopes to unstable or non-remission and how PB T cells and ST pathology are linked, we studied participants from two cohorts of new-onset RA who were treated to a target of remission with csDMARDs, including PB from cohort 1 and paired PB and ST biopsies from cohort 2 (derived from 2 datasets). By studying longitudinal disease activity and cell states from the drug-naïve baseline to 6-12 months after treatment, we identified circulating TIGIT+ Cit-vimentin-autoreactive Tph and CD4+ Tph cells in HLA-DR shared epitope+ RA patients associated with non-remitting RA disease activity in the first year of treatment. These Tph were characterised by lymphoid activation and lymph node egress markers and synovial tissue infiltration with follicular B cell and regulatory T cell (Treg) interaction.

## Research Design and Methods

### Study design and patient characteristics

Cohort 1 was used for hypothesis generation and comprised 21 recent-onset drug-naïve RA participants meeting 2010 ACR/EULAR criteria (12), and carrying HLA-DRB1\**04:01*, \**01:01*, or \**04:04*, who were recruited prospectively to the Australian Arthritis and Autoimmune Biobank Collaborative (A3BC) observational study from the early arthritis outpatient clinic of the Princess Alexandra Hospital in Brisbane, Australia within 3 months of diagnosis. They were treated-to-target of remission with methotrexate, hydroxychloroquine, sulfasalazine, and leflunomide, either singularly or in combination. Disease activity scores using C-reactive protein (DAS28-CRP) and peripheral blood mononuclear cells (PBMCs) were collected at baseline, 6 and 12 months. Stable remission was defined as DAS28-CRP <2.4 at both 6 and 12 months and non-remission was defined as DAS28-CRP ≥2.4 at either 6 months, 12 months or both. This study was approved by the Northern Sydney Local Health District Human Research Ethics Committee (2019/ETH10386) and ratified by the University of Queensland Human Research Ethics Committee (2019001262/HREC/HAWKE/339), with site-specific authorisation obtained via the National Mutual Acceptance (NMA) scheme at Metro South.

To support hypotheses generated in Cohort 1, Cohort 2 comprised 24 recent-onset drug-naïve RA participants meeting 2010 ACR/EULAR criteria, recruited prospectively from the 396.10 study at Flinders Medical Centre, Adelaide, Australia with <12 months’ symptom duration. They were treated to target of remission and scored for DAS28CRP as for cohort 1. Paired PBMCs and arthroscopic ST biopsies were collected at baseline and 6 months post-treatment. Single cell transcriptomic data were collected from CD4+ T cells in 22 participants, and 14 of them were imaged for PD1, CD4, CD20, CD38, THY1, PDPN and DAPI with PhenoCycler spatial proteomics (Akoya). This study was approved by the Southern Adelaide Clinical Human Research Ethics Committee (HREC/20/SAC/3.2), and informed consent was obtained from all participants. To improve statistical power, CD4+ T cell single cell transcriptomic data, from an additional 4 drug-naïve RA participants, analysed previously as part of the Accelerating Medicines Partnership Program: RA and Systemic Lupus Erythematosus (AMP RA/SLE) Network (10), were integrated with the data from cohort 2. Further details are described in Supplementary Methods. A flow chart of participants and disposition of samples in each cohort is shown in Figure 1.

**Figure 1.**
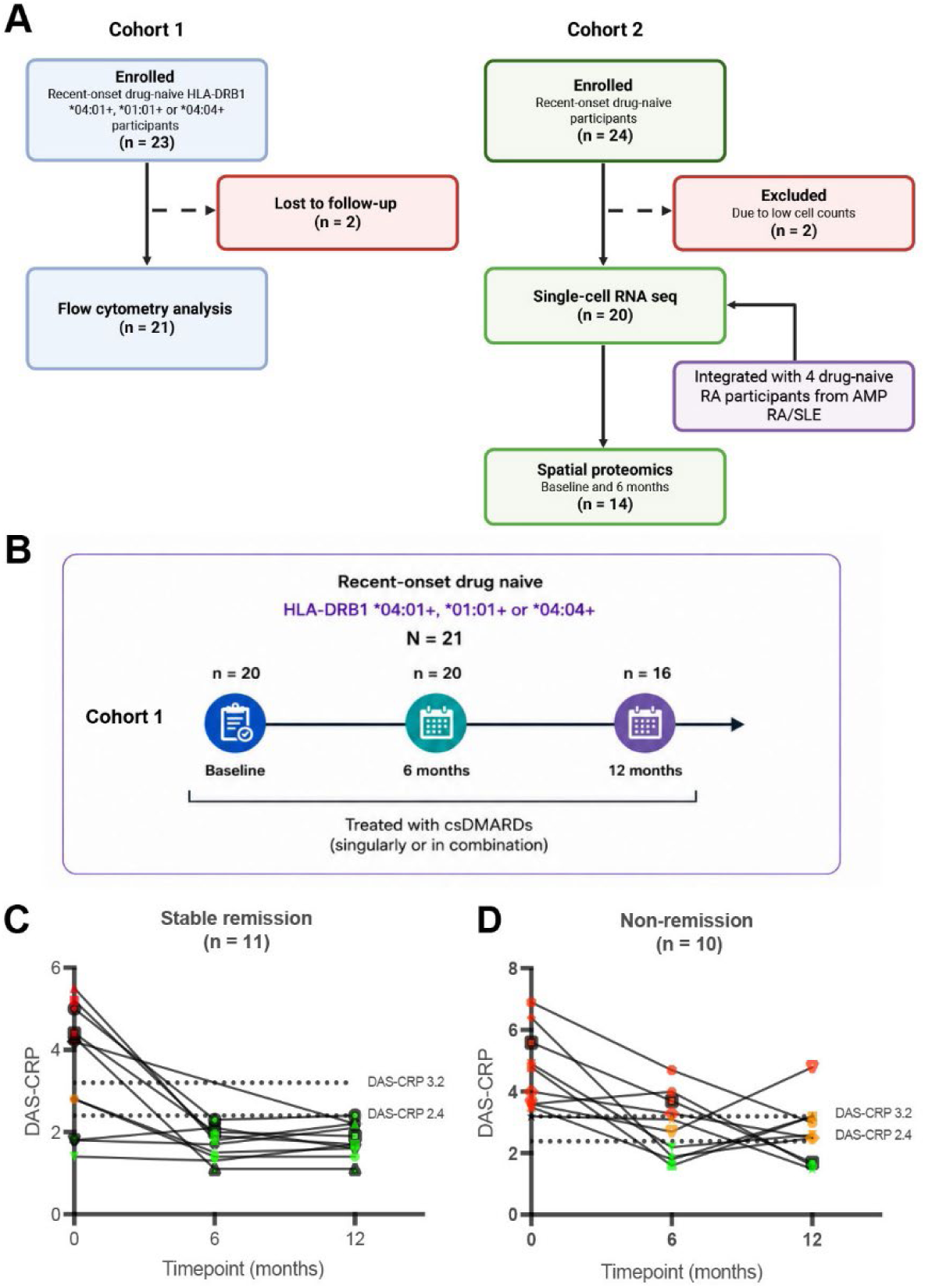
Study design and longitudinal disease activity trajectories in early rheumatoid arthritis following csDMARD treatment. (A) Overall study design of cohorts. (B) Study design for cohort 1. (C-D) Longitudinal DAS28- CRP of drug-naïve early-onset RA participants at baseline, and 6- and 12- months post-treatment with csDMARDs, stratified by stable remission and non-remission. Each line and shape represents an individual participant. Green represents DAS28-CRP <2.4, orange represents DAS28-CRP ≥2.4 – ≤3.2, and red represents DAS28-CRP >3.2.

Patient and public involvement: Patients and consumers have been involved in A3BC since its early development, helping to shape research priorities and biospecimen and data collection strategies to ensure alignment with the needs of people living with musculoskeletal and autoimmune diseases. Consumer input has informed study design, including consent processes, participant information materials, questionnaires, biospecimen collection procedures, and data linkage activities. Patients and consumers continue to provide feedback on study procedures and data collection, ensuring participation remains relevant, meaningful, and acceptable while minimising burden. A3BC also maintains ongoing partnerships with several consumer organisations and regularly includes representatives in the development of grant applications, research projects, and dissemination activities. All patients attending early RA clinics and meeting inclusion criteria were invited to join A3BC or the 396.10 study.

### Spectral flow cytometry and analysis

A 17-marker spectral flow panel incorporated HLA-DRB1*04:01/01:01-VimentinCit64_59-71_ or HLA-DRB1*04:04-VimentinCit71_66-78_ tetramers. If participants tested positive for two of the following HLA alleles: HLA-DRB1\**04:01*, \**01:01*, or *04:04*, priority for tetramer staining was assigned in the following order: HLA-DRB1\**04:01* > *01:01* > *04:04*. Details of cell preparation and gating are in Supplementary Methods, strategy and sample plots are shown in Supplementary Figure 1.

We compared changes in cell population numbers identified with supervised gating over time in remission and non-remission groups using generalized linear mixed modelling with a negative binomial distribution. The fixed effects of timepoint and disease activity status captured temporal changes in cell populations and differences associated with disease activity, respectively, while interaction terms between trajectory and timepoint, and between disease activity and timepoint, assessed whether longitudinal changes differed between stable remission and non-remission groups and across disease activity. Populations of Cit-vimentin-reactive CD4⁺ T cells and total CD4⁺ T cells were analysed separately.

### Generalized linear mixed effect modelling

The numbers of each cell subset over time were modelled using generalised linear mixed modelling with a negative binomial distribution, implemented using the *lme4* package (version 1.1.35.5) in R. Two models, a full and a null model, were fitted. In the full model, the dependent variable was the cell count. Fixed effects include the trajectory (stable remission or non-remission), timepoint (baseline, 6 months, 12 months), clinical outcome (remission, low, moderate/high), and the interaction between trajectory and timepoint, as well as outcome and timepoint. As the other variables were not significantly different between the two groups, we did not adjust for them in the model. Baseline, stable remission and remission disease activity were used as reference categories. A random intercept for each individual (based on a unique subject identifier) was included as a random effect.

In the null model, the terms were identical, except that the fixed effects for trajectory, timepoint, and outcome were excluded. Full models were compared to corresponding null models using an analysis of variance (ANOVA) test to assess the contribution of trajectory, time and outcome to the variation in cell counts, and only statistically significant models were interpreted. An offset term using the logarithm of the parent count of each cell population was included in both models to account for differences in total counts (Supplementary Table 1-4). False discovery rate (FDR) correction was applied using the Benjamini-Hochberg procedure.

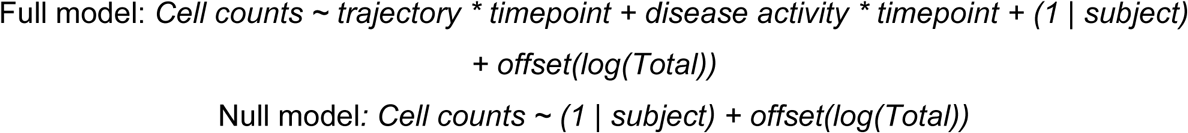

We used a final significance threshold of p<0.05.

### Single-cell transcriptomics and spatial proteomics analysis

Single cells were clustered using Seurat (version 5.4.0), followed by Harmony batch correction. Shared nearest neighbour graph-based Louvain clustering was performed on the Harmony embeddings, and clusters were visualized using uniform manifold approximation and projection (UMAP). Details in Supplementary Methods. Cell-level differential gene expression between TIGIT+ and TIGIT− Tph cells was assessed using the Wilcoxon rank-sum test, with p-values adjusted for multiple testing using the Benjamini–Hochberg method.

Spatial proteomics extraction and preprocessing used the SPACEc workflow (13). Details in Supplementary Methods. Briefly, PhenoCycler images were imported, and tissue regions were extracted based on fluorescence intensity. Cells were segmented using the Cellpose cyto3 model with DAPI as the nuclear guide. Markers included CD79a, CD20, CD21, HLA-DR, CD38, CD138, CD3, CD4, Foxp3, THY1, PDPN and PD1. Cluster identities were assigned based on marker expression profiles (Supplementary Figure 2). Nearest-neighbour distances (µm) between Tfh/Tph cells and Treg, follicular B cells and plasma cells were derived from multiplexed imaging data, in which Delaunay triangulation was constructed from cell centroids, pooling all biopsies. To avoid pseudo-replication, distances were aggregated to a single median value per patient for each cell type, so that each patient contributed one observation per cell type. Because every patient contributes a distance to all three cell types and the distances were non-normally distributed, the three cell types were compared using a Friedman test. Analyses were performed in R using the Tidyverse.

### Protein-protein interaction networks and gene-gene correlation analysis

Protein-protein interaction (PPI) networks were constructed to elucidate interactions between proteins encoded by upregulated genes in CD4+ TIGIT+ Tph and CD4+ TIGIT- Tph cells using Cytoscape (version 3.10.3). A gene-gene Spearman correlation heatmap was generated using patient-level average normalized expression of the top 30 genes upregulated in CD4+ TIGIT+ Tph cells and the top 30 genes upregulated in CD4+ TIGIT- Tph cells.

## Results

### Characteristics of participants

The participant flow diagram detailing enrolment, inclusions and loss to follow-up of cohort 1 is presented in Figure 1A. Of 21 drug-naïve cohort 1 RA participants (Table 1 and Figure 1B), 11 were classified as stable remission (DAS28CRP≤2.4 at 6 and 12 months) and 10 as non-remission (DAS28CRP>2.4 at 6 and/or 12 months). The trajectory of disease activity for each patient is shown in Figure 1C-D. All participants in the stable remission group were ACPA+, and 9 out of 11 were ACPA+RF+. In the non-remission group, 3 out of 10 participants were seronegative. The mean (standard deviation, SD) age of participants was 58.69 (10.70) years in the stable remission group and 58.92 (9.80) years in the non-remission group.

**Table 1.**
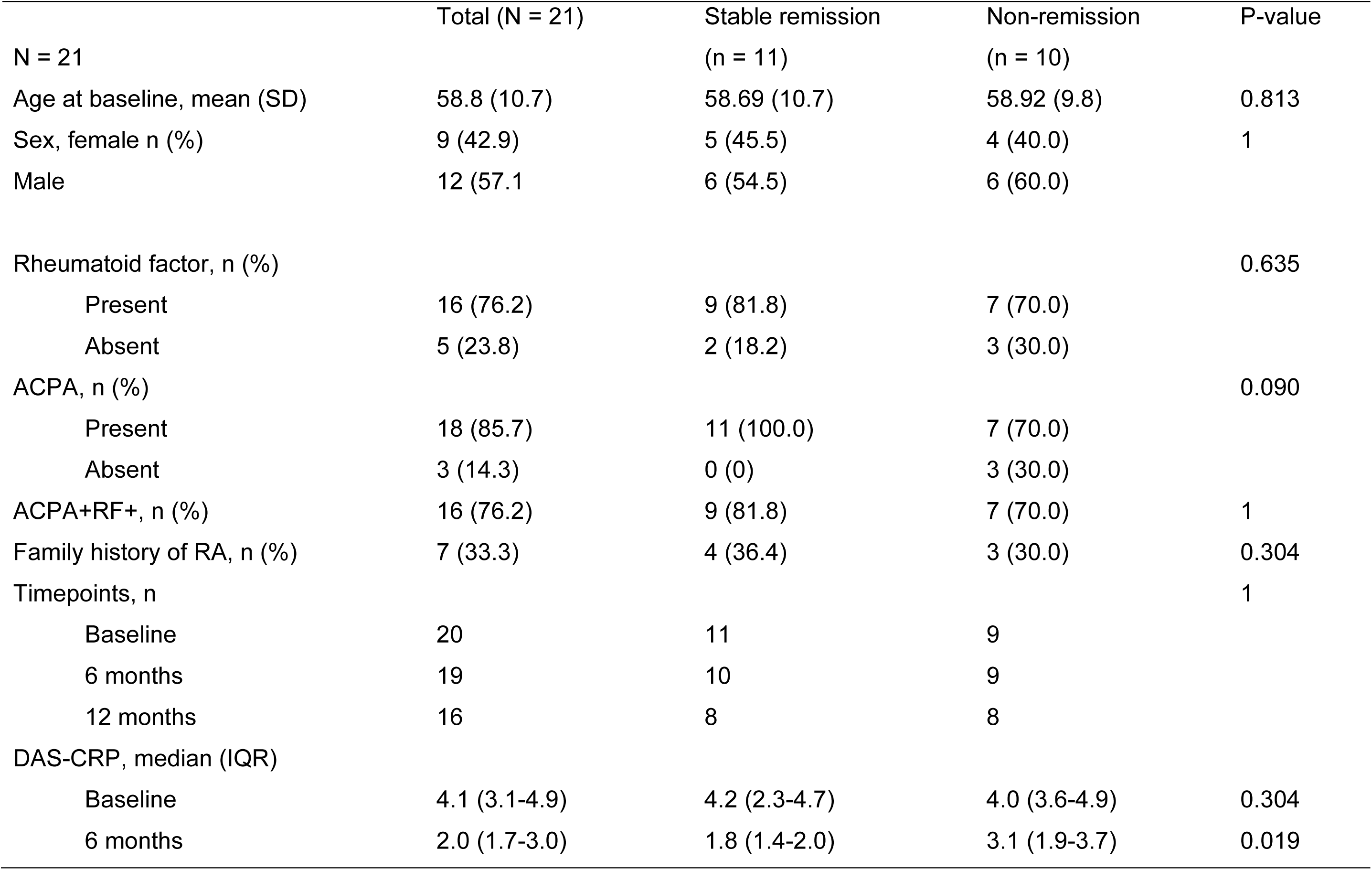
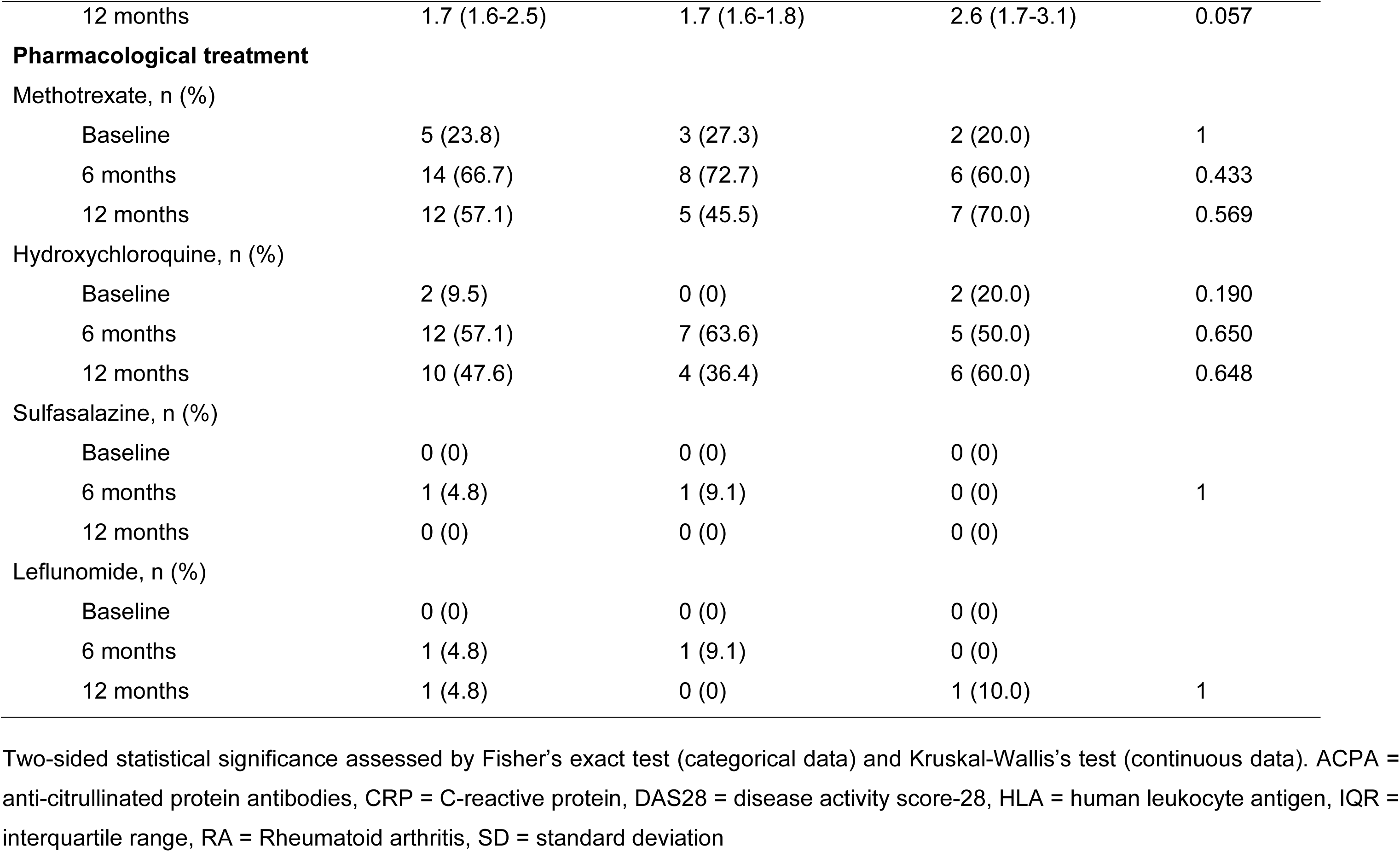
Characteristics of cohort 1 by stable remission and non-remission.

|  | Total (N = 21) | Stable remission<br>(n = 11) | Non-remission<br>(n = 10) | P-value |
| --- | --- | --- | --- | --- |
| N = 21 |  |  |  |  |
| Age at baseline, mean (SD) | 58.8 (10.7) | 58.69 (10.7) | 58.92 (9.8) | 0.813 |
| Sex, female n (%) | 9 (42.9) | 5 (45.5) | 4 (40.0) | 1 |
| Male | 12 (57.1) | 6 (54.5) | 6 (60.0) |  |
| Rheumatoid factor, n (%) |  |  |  | 0.635 |
| Present | 16 (76.2) | 9 (81.8) | 7 (70.0) |  |
| Absent | 5 (23.8) | 2 (18.2) | 3 (30.0) |  |
| ACPA, n (%) |  |  |  | 0.090 |
| Present | 18 (85.7) | 11 (100.0) | 7 (70.0) |  |
| Absent | 3 (14.3) | 0 (0) | 3 (30.0) |  |
| ACPA+RF+, n (%) | 16 (76.2) | 9 (81.8) | 7 (70.0) | 1 |
| Family history of RA, n (%) | 7 (33.3) | 4 (36.4) | 3 (30.0) | 0.304 |
| Timepoints, n |  |  |  | 1 |
| Baseline | 20 | 11 | 9 |  |
| 6 months | 19 | 10 | 9 |  |
| 12 months | 16 | 8 | 8 |  |
| DAS-CRP, median (IQR) |  |  |  |  |
| Baseline | 4.1 (3.1-4.9) | 4.2 (2.3-4.7) | 4.0 (3.6-4.9) | 0.304 |
| 6 months | 2.0 (1.7-3.0) | 1.8 (1.4-2.0) | 3.1 (1.9-3.7) | 0.019 |
| 12 months | 1.7 (1.6-2.5) | 1.7 (1.6-1.8) | 2.6 (1.7-3.1) | 0.057 |
| **Pharmacological treatment** |  |  |  |  |
| Methotrexate, n (%) |  |  |  |  |
| Baseline | 5 (23.8) | 3 (27.3) | 2 (20.0) | 1 |
| 6 months | 14 (66.7) | 8 (72.7) | 6 (60.0) | 0.433 |
| 12 months | 12 (57.1) | 5 (45.5) | 7 (70.0) | 0.569 |
| Hydroxychloroquine, n (%) |  |  |  |  |
| Baseline | 2 (9.5) | 0 (0) | 2 (20.0) | 0.190 |
| 6 months | 12 (57.1) | 7 (63.6) | 5 (50.0) | 0.650 |
| 12 months | 10 (47.6) | 4 (36.4) | 6 (60.0) | 0.648 |
| Sulfasalazine, n (%) |  |  |  |  |
| Baseline | 0 (0) | 0 (0) | 0 (0) |  |
| 6 months | 1 (4.8) | 1 (9.1) | 0 (0) | 1 |
| 12 months | 0 (0) | 0 (0) | 0 (0) |  |
| Leflunomide, n (%) |  |  |  |  |
| Baseline | 0 (0) | 0 (0) | 0 (0) |  |
| 6 months | 1 (4.8) | 1 (9.1) | 0 (0) |  |
| 12 months | 1 (4.8) | 0 (0) | 1 (10.0) | 1 |

Two-sided statistical significance assessed by Fisher's exact test (categorical data) and Kruskal-Wallis's test (continuous data). ACPA = anti-citrullinated protein antibodies, CRP = C-reactive protein, DAS28 = disease activity score-28, HLA = human leukocyte antigen, IQR = interquartile range, RA = Rheumatoid arthritis, SD = standard deviation

DAS28-CRP scores did not differ significantly between the groups at baseline, but both groups improved with treatment, as DAS28-CRP scores decreased significantly at 6 months and 12 months for each of the stable remission (p = 0.006) and non-remission (p = 0.007) groups. DAS28-CRP scores were significantly lower in the stable remission group compared to the non-remission group at 6 months (p = 0.019) and trended lower at 12 months (p = 0.057). There were no significant differences in DAS28-CRP scores between ACPA+ and ACPA- participants at baseline, 6 months or 12 months.

### Moderate/high disease activity is associated with higher CD4+ Tph frequencies

Cit-vimentin-reactive CD4+ T cells were characterised in PBMCs collected from all donors at all time points using either HLA-DRB1*04:01/01:01–Vimentin_59-71_Cit64 or HLA- DRB1*04:04–Vimentin_66-78_Cit71 tetramers. Twenty-six populations were identified with supervised gating.

Among the 26 CD4⁺ and 26 Cit-vimentin-reactive CD4+ T cell populations assessed with supervised gating, in moderate/high disease activity multiple CD4+CD45RA-CXCR5-PD1+ non-regulatory T cell populations characteristic of Tph increased, as compared to remission observations, regardless of timepoint or trajectory (Figure 2A-F). These included CD4+ PD1+TIGIT+ (p = 0.018), TIGIT+ (p = 0.018), PD1+ Tph (p = 0.018), CCR7+PD1+ stem-like Tph (p <0.001). Cit-vimentin-reactive CD4+CD45RA-CXCR5-PD1+ Tph (p = 0.043) and Cit-vimentin-reactive CD25^hi^CD127^-^ Treg (p = 0.043) were also increased in moderate/high disease activity. These findings remained unchanged with exclusion of seronegative participants, with the exception of Cit-vimentin-reactive Tph cells, which did not reach statistical significance (p = 0.052).

**Figure 2.**
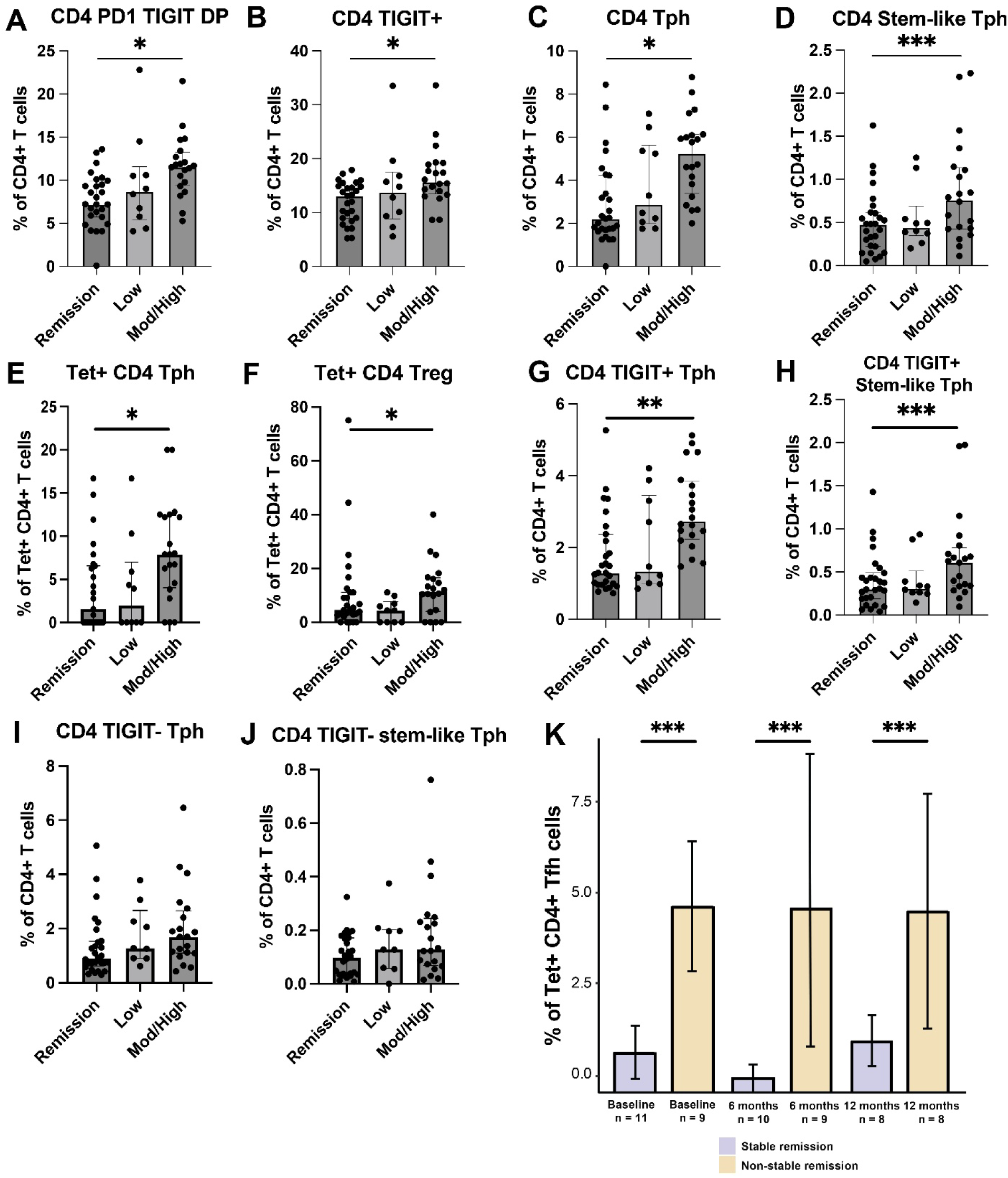
Disease activity and treatment response are associated with changes in circulating CD4+ PD1+ T cell subsets. (A-H) Bar plots demonstrating the frequency of differentially expressed CD4+ T cells clustered using manual (supervised) gating stratified by disease severity. Low = Low disease activity at point of observation, Mod/High = Moderate or high disease activity at point of observation, Remission = Remission at point of observation. (K) Bar plots of differentially expressed supervised gated citrullinated-vimentin-specific CD4^+^ Tfh cells over time (baseline, 6 months and 12 months) stratified by stable remission and non-remission.

Given the similar trends observed in CD4⁺ PD1⁺TIGIT⁺ and TIGIT⁺ T cells, we explored their overlap using Boolean and flow gating. In PB, TIGIT+ Tph are a subset of Tph expressing high levels of PD1. The CCR7+ stem-like Tph subset expresses higher cell surface TIGIT and lower PD1 than the CCR7- Tph (Supplementary Figure 3). We determined that this TIGIT+ subset drives the association of Tph with non-remission, as TIGIT+ Tph (p = 0.002) and TIGIT+ stem-like Tph (p < 0.001), but neither TIGIT- Tph nor TIGIT- stem-like Tph were increased in moderate/high disease activity observations, as compared to remission observations, regardless of timepoint or trajectory (Figure 2G-H).

### Lower citrullinated-vimentin-reactive CD4+ Tfh post-treatment associated with stable remission

Immune trajectories of Cit-vimentin-reactive follicular T helper cells (Tfh) were increased in non-remission participants, as compared to stable remission participants, over time from baseline to 6 months (p = <0.001) (Figure 2I).

At 6 months post-treatment, higher frequencies of Cit-vimentin-reactive Tfh cells identified through supervised gating, were significantly associated with non-remission compared with stable remission (negative binomial regression, β = 3.65, p = 0.042). In contrast, baseline frequencies of Cit-vimentin-reactive Tfh cells were not significantly associated with disease trajectory (p = 0.135). These findings remained unchanged with the exclusion of seronegative participants.

### Unsupervised clustering reveals CD25^+^CD127^+^ Tem expressing either CCR6 or CCR4 decrease by 6 months when disease remains active on treatment

To complement and extend the manually gated differentially expressed populations, unsupervised clustering using FlowSOM was applied to leverage high-dimensional marker data and identify cell populations in an unbiased manner by integrating all available markers. Fifteen CD4⁺ and 14 Cit-vimentin-specific CD4⁺ T cell clusters were identified from 30 meta- clusters (Figure 3A-C), including 4 clusters of Tph and Tfh, based on marker expression (Figure 3D).

**Figure 3.**
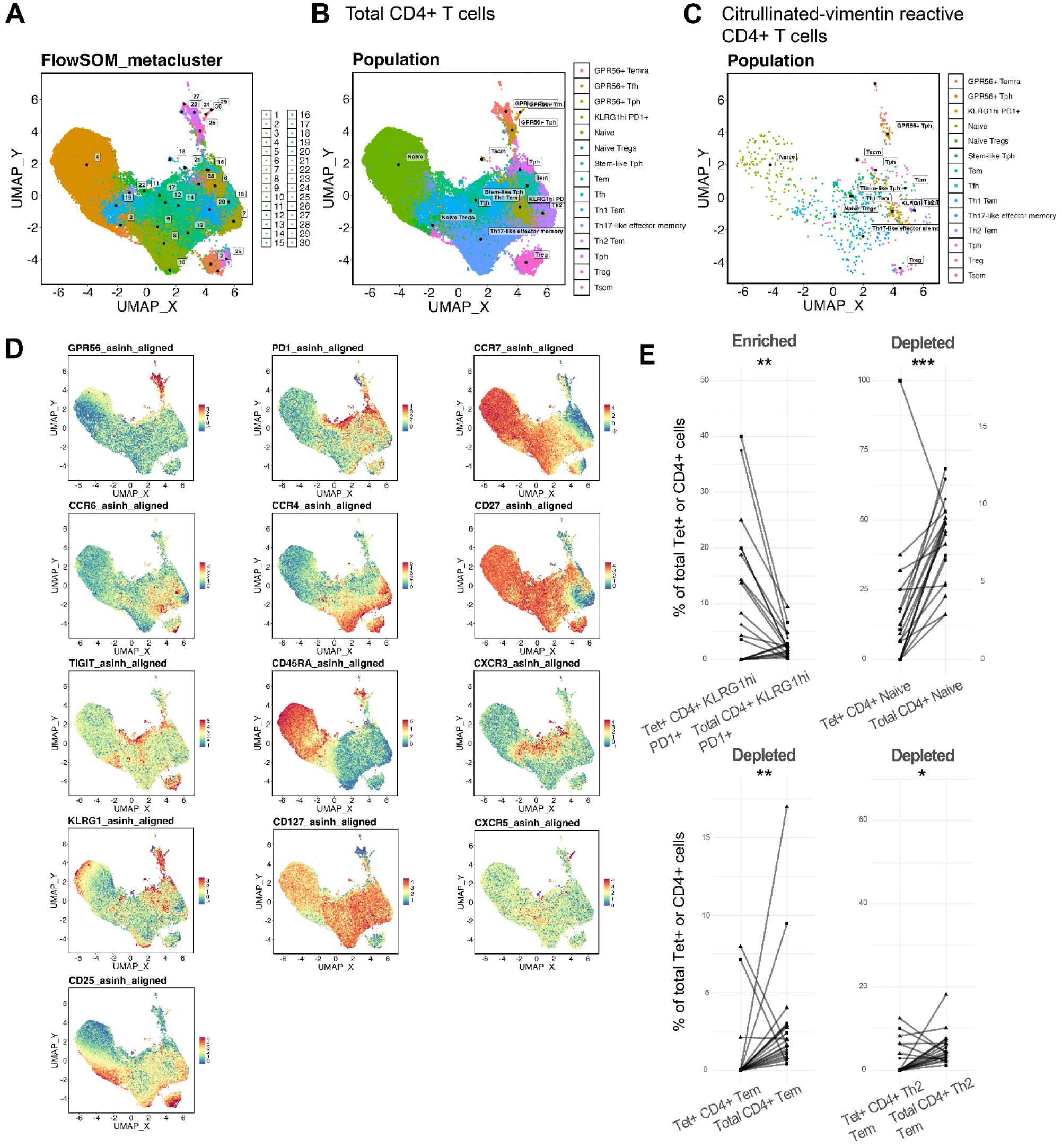
Unsupervised characterization of CD4^+^ T-cell subsets. FlowSOM clustering after dimensionality reduction plotted using UMAP, of (A) CD4^+^ T cells FlowSOM meta-clusters, (B) annotated CD4^+^ T cells and (C) annotated Cit-vimentin specific CD4^+^ T cells, with (D) multi-plots demonstrating individual marker expression across cells. (E) Enrichment plots comparing the percentages of Cit-vimentin-reactive CD4+ T cells and CD4+ T cells from unsupervised clustering that were significantly enriched or depleted at baseline. Each line connects the frequency of the indicated subset among Cit-vimentin- reactive CD4+ T cells and among total CD4+ T cells from the same participant. Paired comparison was performed using the Wilcoxon signed rank test.

At baseline, Cit-vimentin-specific CD4^+^ T cells were significantly enriched in a KLRG1^hi^PD1^+^CCR6^+^CXCR3^lo^ antigen-experienced subset relative to total CD4^+^ T cells (Figure 3E). Conversely, at baseline, Cit-vimentin-specific CD4^+^ T cells were significantly depleted in naïve, CD45RA^-^CD25^+^CD127^+^CCR6^+^ T effector memory (Tem) and a CD45RA^-^ CCR4^+^CD25^+^CD127^+^ T helper-2 effector memory (Th2 Tem) subset, previously described to be increased in partial remission in type 1 diabetes (14).

Between baseline and 6 months, CD4⁺ Tem (p = 0.043) and CD4⁺ Th2 Tem (p = 0.043) trajectories were significantly more likely to decrease in visits classified as moderate/high disease activity (DAS28-CRP>2.4) compared with remission visits, during which trajectories increased (Figure 4). Between 6 and 12 months these trends continued but were no longer statistically significant. These data indicate that CD25^+^CD127^+^ Tem expressing either CCR6 or CCR4 change in a disease activity-dependent manner, particularly in the first months of treatment.

**Figure 4.**
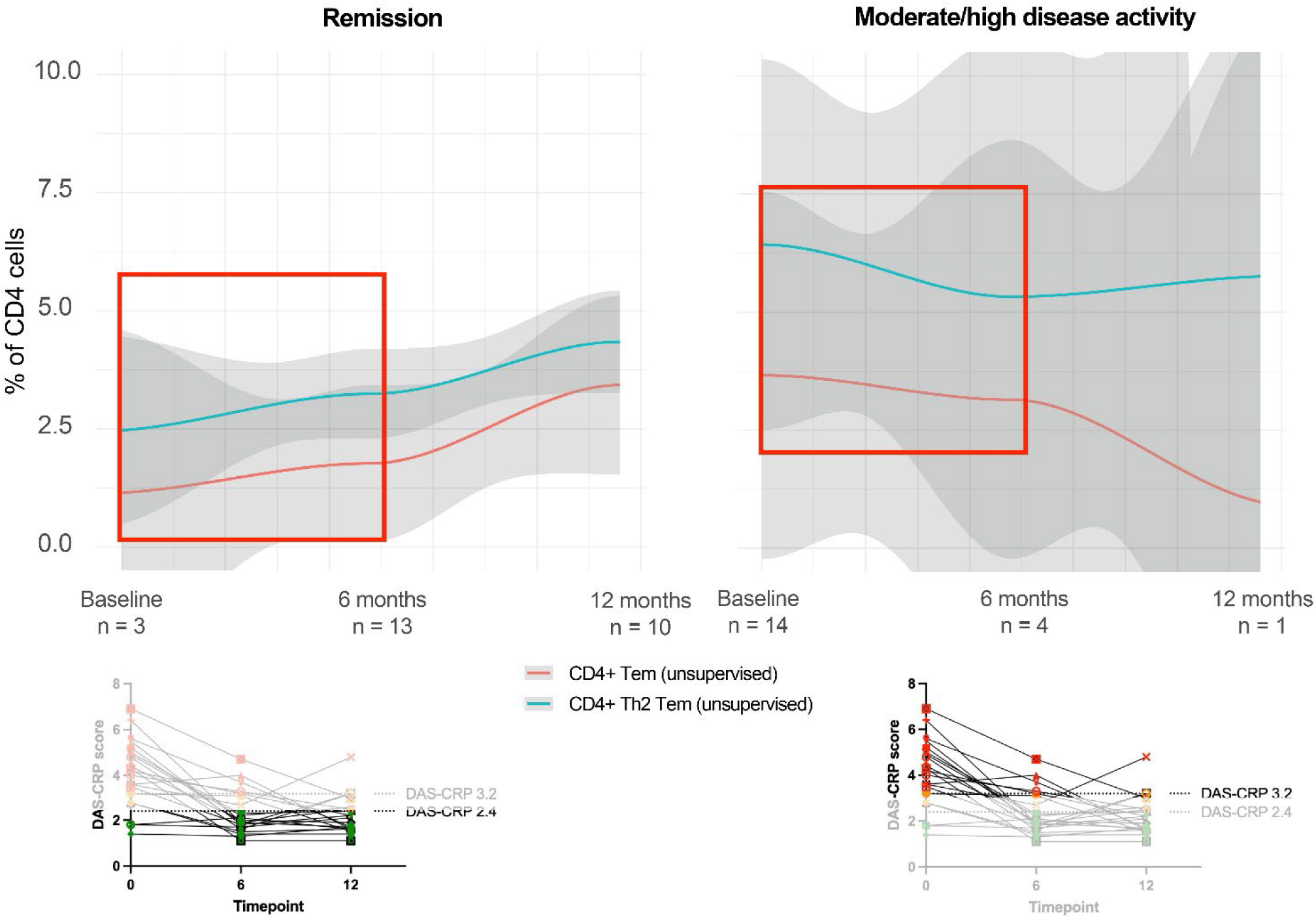
Longitudinal modelling of CD4+ T-cell populations by disease activity. Spline trajectories of unsupervised clustered CD4^+^ Tem and CD4^+^ Th2 cells over time (baseline, 6 months and 12 months) stratified by moderate/high and remission disease activity. Grey shaded areas represent the 95% confidence interval. Red boxes identify significant timepoints. Dot plots demonstrate DAS28-CRP of RA participants in (left) moderate/high disease activity or (right) remission over time. Green represents DAS28-CRP <2.4, orange represents DAS28-CRP ≥2.4 – ≤3.2, and red represents DAS28-CRP >3.2.

While non-significant, the trends for CD4+ PD1+ populations (CD4+ Tph, CD4+ stem-like Tph, and CD4+ Tfh) from unsupervised clustering were similarly increased in moderate/high disease activity as observed in supervised gating (Supplementary Figure 4A-B), and immune trajectories of CD4⁺ Tfh were increased in non-remission participants as compared to stable remission participants (Supplementary Figure 4C).

### Remodelling of the circulating Tph compartment identified by trajectory-specific longitudinal modelling

Next, we investigated longitudinal changes within stable remission and non-remission participants. In participants achieving stable remission, several CD4⁺ T cell subsets changed significantly over time (Table 2). At 12 months, there was a reduction in CD4⁺ stem-like Tph, CD4⁺ TIGIT⁺ stem-like Tph, CD4⁺ TIGIT⁺ Tph, CD4⁺ TIGIT⁺, and CD4⁺ KLRG1⁺ populations. Conversely, CD4⁺ CXCR3⁺CCR4⁺ cells and CD4⁺ CXCR3⁺CCR4⁻ cells increased at 12 months. From unsupervised clustering, a decrease in CD4⁺ Tph was evident at 6 months.

**Table 2.**
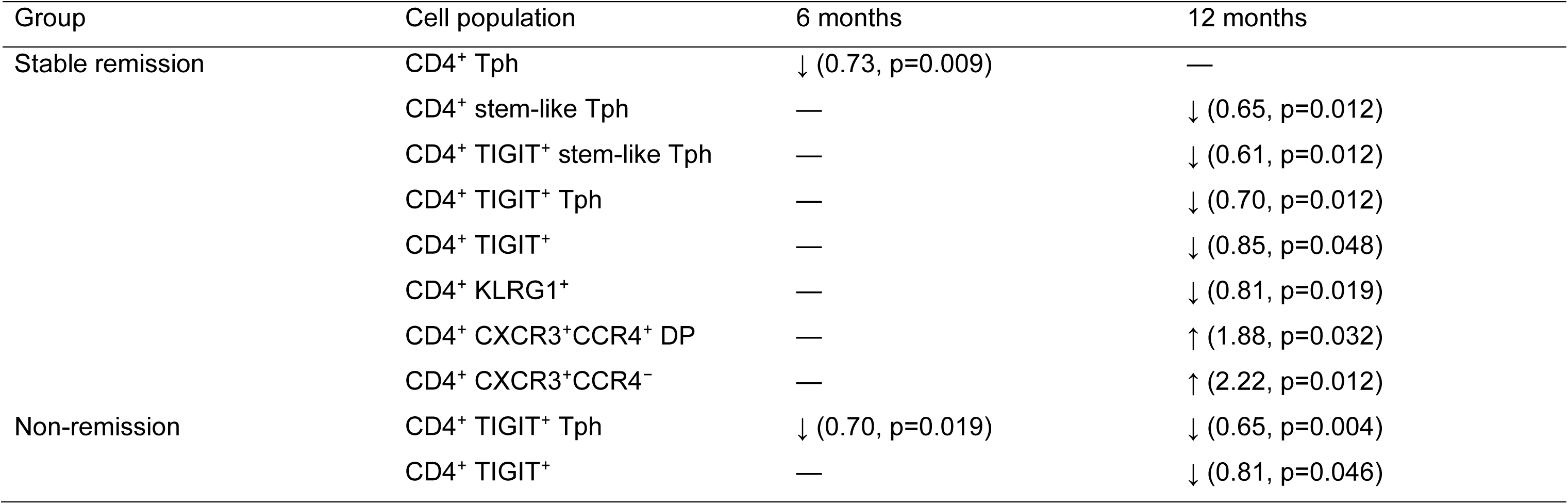
Changes in T cell subsets in stable remission and non-remission RA participants using Incidence Rate Ratio.

| Group | Cell population | 6 months | 12 months |
| --- | --- | --- | --- |
| Stable remission | CD4<sup>+</sup> Tph | ↓ (0.73, p=0.009) | — |
|  | CD4<sup>+</sup> stem-like Tph | — | ↓ (0.65, p=0.012) |
|  | CD4<sup>+</sup> TIGIT<sup>+</sup> stem-like Tph | — | ↓ (0.61, p=0.012) |
|  | CD4<sup>+</sup> TIGIT<sup>+</sup> Tph | — | ↓ (0.70, p=0.012) |
|  | CD4<sup>+</sup> TIGIT<sup>+</sup> | — | ↓ (0.85, p=0.048) |
|  | CD4<sup>+</sup> KLRG1<sup>+</sup> | — | ↓ (0.81, p=0.019) |
|  | CD4<sup>+</sup> CXCR3<sup>+</sup>CCR4<sup>+</sup> DP | — | ↑ (1.88, p=0.032) |
|  | CD4<sup>+</sup> CXCR3<sup>+</sup>CCR4<sup>-</sup> | — | ↑ (2.22, p=0.012) |
| Non-remission | CD4<sup>+</sup> TIGIT<sup>+</sup> Tph | ↓ (0.70, p=0.019) | ↓ (0.65, p=0.004) |
|  | CD4<sup>+</sup> TIGIT<sup>+</sup> | — | ↓ (0.81, p=0.046) |

In non-remission participants, CD4⁺ TIGIT⁺ Tph decreased at both 6 months and 12 months, alongside a reduction in CD4⁺ TIGIT⁺ cells at 12 months. These data indicate that while higher Tph frequencies are associated with higher disease activity overall, immune remodelling involving circulating Tph occurs over the first 12 months, as RA is treated and disease activity improves.

### Lower CD4+PD1+ T cells associated with follicular B cells in synovial tissue in remission than non-remission

The ongoing high levels of circulating Cit-vimentin-specific Tfh in non-remission suggested persistent T-B interactions in lymphoid tissue. Furthermore, as Tph remodel in blood with treatment, it is important to understand whether this process is also occurring in ST. We therefore compared PhenoCycler spatial proteomics analysis of ST biopsies of cohort 2 participants in remission or non-remission after 6 months’ treatment with DMARDs. We identified CD79a+CD20+CD21+HLA-DR+ B cells located in follicular aggregates, CD79a+CD20+CD21+CD38+CD138+ plasma cells, CD79a+CD38+ plasmablasts, CD3+CD4+Foxp3+ Treg, THY1+ pericytes surrounding blood vessels and PDPN+ fibroblasts and CD4+PD1+ Tph/Tfh cells.

From the baseline (DAS28-CRP = 5.71) and 6 months (DAS28-CRP = 4.94) biopsies of a representative non-remission participant, CD4+PD1+ T cells localise predominantly to follicular aggregates, within T cell zones adjacent to follicular B cells. Plasmablasts and plasma cells were located extra-follicularly (Figure 5A). In contrast, in the representative remission biopsy (DAS28-CRP = 2.15), follicular lymphoid aggregates were much smaller, and plasmablasts/plasma cells much fewer as compared to baseline (DAS28-CRP = 6.48) (Figure 5B). The number of ST infiltrating Tph/fh cells correlated with the number of infiltrating Treg (R^2^ 0.963, p<0.0001), follicular B cells (R^2^ 0.966, p<0.0001), plasmablasts (R^2^ 0.584, p=0.0015) and plasma cells (R^2^ 0.462, p=0.0075), consistent with inflammatory co-recruitment. Spatial proximity calculations in pre-treatment biopsies (n=12) identified Tph/fh nearest to Treg, with follicular B cells at a similar distance, and plasma cells significantly more distant (p = 0.017, Figure 5C).

**Figure 5.**
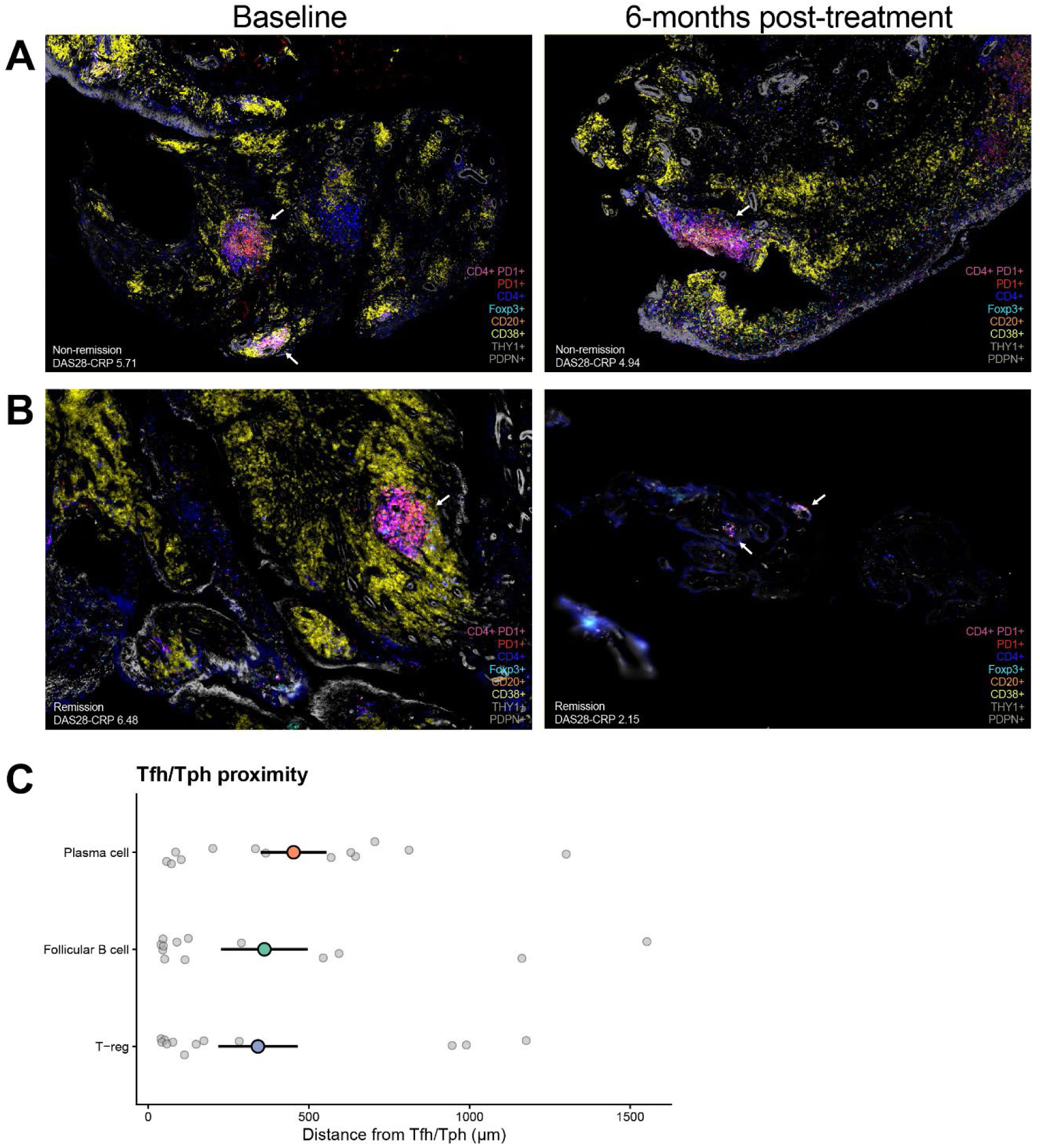
Spatial distribution of CD4+ PD1+ T cells, B cells and plasma cells in synovial tissue before and after csDMARD treatment. Spatial proteomics (PhenoCycler) images of synovial tissue (magnification: 200μm) showing CD4+ Tfh/Tph cells (stained with CD4 and PD1 in magenta), PD1+ cells (PD1 in red), CD4+ cells (CD4 in dark blue), B cells (CD20 in orange), plasma cells (CD38 in yellow), regulatory T cells (Foxp3 in cyan), peri-vascular pericytes (THY1 in grey), and fibroblast (podoplanin in grey) in ACPA+ RA patients who (A) did not and (B) did achieve DAS28 remission on conventional synthetic disease-modifying antirheumatic drugs at baseline and 6 months post-treatment. DAS28-CRP = Disease Activity Score in 28 joints using C-reactive protein. C: Spatial proximity of Tfh/Tph cells to Treg, follicular B cells and plasma cells across all biopsies combined. Each grey dot is one patient’s median nearest-neighbour distance from Tfh/Tph to the indicated cell type; coloured points show the group mean ± SEM. Plasma cells were significantly further than Treg or follicular B cells from Tfh/Tph (p=0.017, Friedman test).

### TIGIT+ Tph are transcriptionally activated for immune responses

Since circulating TIGIT+ and Cit-vimentin-reactive Tph and Tph/fh cells associated with ST follicular B cells were strongly associated with disease activity in early RA, we analysed a single-cell RNAseq dataset comprising PB and ST CD4+ T cells from cohort 2 and additional data from the previously published AMP RA/SLE consortium, for differentially expressed genes characterising CD4⁺ Tph in PB and ST (Supplementary Table 5). Eighteen participants (396.10 study, n = 15; AMP RA/SLE, n = 3) were ACPA+. Nine CD4+ UMAP clusters were identified (Figure 6A and Supplementary Figure 5), of which the Tph/Tfh compartment expressing *PD1* and *CXCL13* could be further separated into proliferating, stem-like and *TIGIT*-expressing effector-like Tph/Tfh populations (Figure 6B). Effector-like and proliferating Tph/Tfh were enriched in ST while stem-like Tph/Tfh were enriched in PB (Figure 6C).

**Figure 6.**
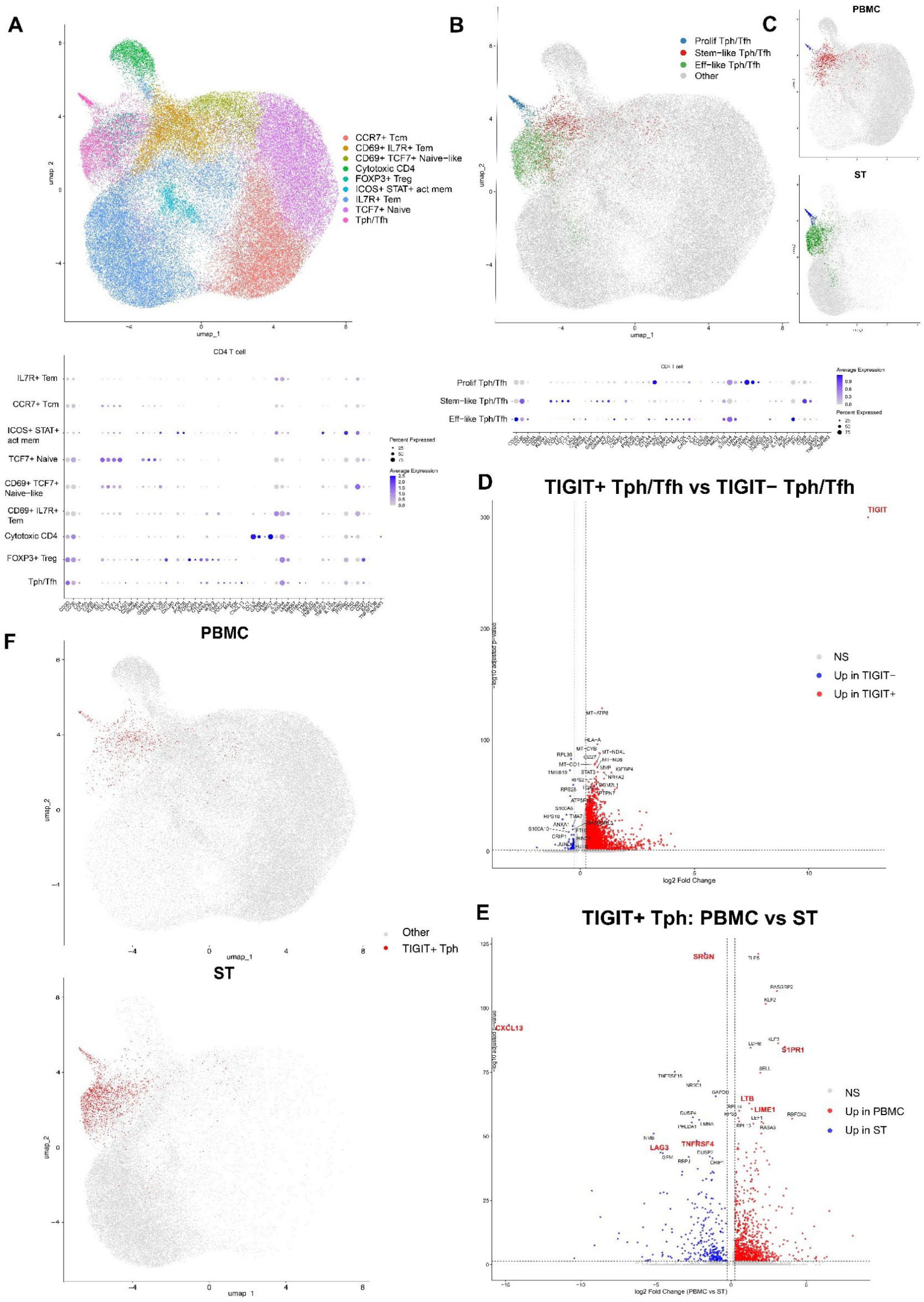
Single-cell transcriptomic characterization of CD4+ Tph cell populations in peripheral blood and synovial tissue. UMAP of scRNA-seq data for (A) CD4^+^ T cells from PBMC and ST and (B) CD4^+^ Tph subsets with dot plots of genes used for cluster annotation. Colour and size of the dots indicate average expression and percent expressed. C: Distribution of CD4^+^ Tph subsets across ST and PB compartments. D: Volcano plot of genes differentially expressed between CD4⁺TIGIT⁺ (red, adjusted p-value < 0.05, log₂FC > 0.25) and TIGIT⁻ Tph/Tfh cells (blue, adjusted p-value < 0.05, log₂FC < −0.25). Non-significant genes are shown in grey. TIGIT is highlighted in red. E: Volcano plot of genes differentially expressed between CD4⁺ TIGIT⁺ Tph/Tfh cells in PB and ST. Genes significantly upregulated in TIGIT⁺ Tph cells in PB (red, adjusted p-value < 0.05, log₂FC > 0.25) and in ST are shown in blue (adjusted p-value < 0.05, log₂FC < −0.25). F: Distribution of TIGIT⁺ Tph cells across ST and PB compartments.

We uncovered marked transcriptional differences between CD4⁺ TIGIT⁺ Tph and CD4⁺ TIGIT⁻ Tph cells, with the majority of differentially expressed genes (326 out of 386) upregulated in the TIGIT⁺ subset (Figure 6D and Supplementary Table 6), suggesting that CD4⁺ TIGIT⁺ Tph are more transcriptionally activated than exhausted. TIGIT+ and TIGIT− Tph cells were of similar quality, with comparable total unique molecular identifier counts and mitochondrial transcript percentages. Ribosomal genes were lower in TIGIT+ Tph as compared to TIGIT− Tph. The increased number of detected genes in TIGIT+ Tph cells is more likely to reflect transcriptional activity associated with activation than differences in sequencing quality or RNA capture efficiency (Supplementary Figure 6).

Gene-gene correlation of genes differentially expressed between TIGIT⁺ and TIGIT⁻ Tph/Tfh cells demonstrates that genes upregulated in CD4⁺ TIGIT⁺ Tph cells are strongly co-expressed and form a large coordinated transcriptional module that is mainly enriched for immune system processing and immune response, suggesting a shared activation-associated program, whereas genes upregulated in CD4⁺ TIGIT⁻ Tph cells have weaker gene-gene co-expression and form more fragmented transcriptional modules that are mainly enriched for peptide metabolic processes (Supplementary Figure 7A and B). Comparison of CD4⁺ TIGIT⁺ Tph cells in PB and ST identified genes associated with T cell receptor-B cell receptor signalling (*LIME1),* lymphoid follicle development *(LTB)* and lymph node egress (*S1PR1)* in circulating TIGIT+ Tph, consistent with recent interaction with antigen presenting B cells in lymphoid organs. On the other hand, CD4⁺ TIGIT⁺ Tph in ST differentially expressed functional B cell chemotactic (*CXCL13*) granule storage (*SRGN*) and regulatory/anergic genes (*LAG3, TNFRSF18)* (GITR), consistent with their location adjacent to follicular B cells and Treg, respectively, in ST aggregates (Figure 6E and 6F).

## Discussion

In this study, we investigated changes in PB total and Cit-vimentin-autoreactive T cells from recent-onset seropositive and seronegative HLA-DR-SE+ RA participants treated over 12 months to a target of remission with csDMARDs. Moderate and high disease activity were associated with high levels of CD4⁺ Tph, including Cit-vimentin-reactive CD4⁺ Tph expressing TIGIT, and non-remission was associated with an increase in CD4⁺ Tfh cells over time. Supporting these findings, non-remission in a second cohort was characterised by large ST follicular aggregates in which Treg and B cells colocalised with CD4+PD1+ Tph/Tfh, surrounded by plasma cells and plasmablasts, whereas in remission, follicular aggregates were sparse, much smaller and colocalised Treg and B cells were further away from Tph/Tfh. This suggests that the enrichment of antigen-experienced PD1+ flow cytometric phenotypes that we observed among Cit-vimentin CD4⁺ T cells in early HLA-DR- SE+ RA reflects ongoing antigen presentation, and *in vivo* activation and differentiation. Indeed, transcriptional analysis demonstrated that circulating CD4⁺ TIGIT⁺ Tph cells express *LIME1, LTB* and *S1PR1* associated with B cell activation, lymphoid expansion and lymph node exit, suggesting that high disease activity reflects the transit of these immune activating T cells from lymph node to ST, where they represent a functionally distinct tissue effector subset recruited in active disease.

CD4^+^CCR7+ stem-like Tph expressing TIGIT were most closely associated with increased disease activity in our flow cytometric study. Stem-like Tph cells were reported in PB and within tertiary lymphoid structures in the synovium, where they colocalised with B cells but were capable of self-renewal and differentiation into effector-like Tph (11). Tfh and Tph support B-cell maturation and production of autoantibodies in secondary and ectopic lymphoid tissue (15, 16) and ex vivo (7). Autoreactive ACPA+ B cells in PB similarly expressed T cell costimulatory ligands, proinflammatory cytokines and could secrete neutrophil chemokines in response to citrullinated antigens, including vimentin. This state of activation was shown to be at its peak in early RA (17).

CD11c+Tbet+ autoimmune-associated B cells (ABCs) that were increased in the circulation and correlated with bone destruction in early RA were antigen presenting cells that differentiated CD4+ T cells to Th17 *ex vivo* (18), a phenotype resembling the antigen-experienced KLRG1^+^PD1^+^CCR6^+^ Cit-Vimentin-reactive T-cell enriched phenotype in the current studies and in cancer (19). Thus, our data support the conclusion that Cit-vimentin- reactive CD4+ T cells in active early RA are responding to antigen presentation by multiple APCs, including autoreactive B cells as well as ABCs, which may have phagocytised and processed Cit-vimentin epitopes (20). This conclusion is supported by previous studies that found the frequency of PB Tph and TIGIT^+^ Tph correlates with disease activity in early RA (21–23) and that CD4⁺ PD1⁺ T cells are expanded in PB and differentiated in the synovium of seropositive and seronegative individuals with RA, consistent with sustained synovial antigen response (7, 23–25).

The transcriptional profile of TIGIT+ Tph cells suggests that TIGIT marks a highly activated, antigen-experienced Tph subset in ST. Although TIGIT can function as an inhibitory receptor, it is also upregulated in response to chronic antigen stimulation, thus can identify highly differentiated effector populations (26). Given the differential expression of their B cell activation, lymphoid expansion and lymph node exit genes, it is notable that CD4⁺ TIGIT⁺ T cells were increased in PB of seropositive RA first-degree relatives of Indigenous North Americans, who have strong enrichment of HLA-SE alleles with capacity to present Cit-vimentin (2, 22).

We found higher levels of Cit-vimentin-reactive CD4^+^ Tfh were associated with a non-remission disease trajectory, and by 6 months this difference distinguished the trajectories. This implies B-cell driven Cit-vimentin antigen presentation to Tfh is sustained in cohort 1 participants who did not achieve remission. Consistent with this, spatial proteomics demonstrated that non-remission cohort 2 participants had ongoing infiltration and colocalization of CD4^+^ PD1^+^ T cells, Treg and follicular B cells in lymphoid aggregates, as compared to remission ST. Although their lineage relationship is not clear yet, the colocalisation of Tph/fh with Treg suggests potential for differentiation from Tph/fh to follicular-regulatory T cells (Tfr) (27, 28). Together the PB and ST data suggest traffic of TIGIT+ Tph from lymph nodes to ST, CXCL13-mediated B cell recruitment to ST. Here, ongoing synovial antigen presentation may support the persistence, and induction of anergy genes in antigen-specific Tfh and Tph. Although expanded PD1^+^ T cells, plasma cells and ABCs have been shown in ST of RA patients by immunofluorescent staining (7, 18), these studies did not link disease activity. Our data clearly show that in remission low circulating levels of Tph reflect low tissue infiltration and not simply migration from PB to ST. This is consistent with reduced chemoattraction of Tph via CCR5 or CCR2 in the face of low synovial disease activity, and in turn reduced tissue-localized T-B cell interactions due to reduced Tph-mediated production of CXCL13 and IL-21 that promote recruitment and engagement of Tfh and B cells (7).

FlowSOM clustering independent of predefined gating strategies identified additional disease activity-dependent decline in CD4^+^ CD25+CD127+CCR6+ Tem and CD4^+^ CD25+CD127+CCR4+ Th2 Tem in moderate and high disease observations. Th2 Tem have been described in children and adolescents with type 1 diabetes associated with longer duration of partial remission after onset. Th2 Tem displayed clonal expansion and expressed IL-4 and IL-10, with anti-inflammatory function (14). Thus, the decrease of Th2 Tem in moderate/high disease activity may reflect RA-associated suppression of Th2-associated immune-regulatory responses, potentially due to the dominance of Th1 over Th2 in peripheral blood (29).

Our analysis of cohort 1 was limited by a relatively small sample size. Furthermore, although both cohorts included seronegative participants, the numbers were small (cohort 1, n = 3, cohort 2, n = 8). However, our study is strengthened by multiple longitudinal time points for cohort 1 and our capacity to strengthen hypotheses generated in cohort 1 with an additional cohort comprised of datasets from 2 separate cohorts. Sensitivity analyses excluding seronegative cohort 1 participants demonstrated that the differentially expressed PB populations identified by flow cytometry remained significant between the specified groups, even after FDR correction. This suggested that for RA patients carrying HLA-DR-SE, the associations of disease activity with Tph, Tfh and Cit-vimentin reactivity did not depend on presence of ACPA. Consistent with this, in cohort 2, we found similar ST infiltration by CD4+PD1+ T cells into B cell and plasma cell rich lymphoid aggregates in seropositive and seronegative RA and similar reductions in remission. It will be interesting to determine whether different antigen presenting B cells might stimulate autoreactive Tph in seronegative RA.

Taken together, our data identify persistently high circulating TIGIT^+^ Tph and Tfh, including Cit-vimentin specificities, that reflect Tph antigen-presenting B-cell interactions in lymph nodes and ST, associate with reduced response to csDMARDs in recent-onset RA. These findings suggest the potential of TIGIT+ Tph blood monitoring to assess requirement for treatment escalation during treat-to-target monitoring.

## Acknowledgements and affiliations

Acknowledgements

The A3BC team are sincerely grateful for the generosity of the patient participants, carers, clinicians, consumer groups and researchers in continuing to give their time and effort to allow research discovery such as that herein. We acknowledge support by the Accelerating Medicines Partnership Program: Rheumatoid Arthritis and Systemic Lupus Erythematosus (AMP RA/SLE) Network for data used in the analysis.

## Author contribution

J.H. contributed to writing—original draft preparation, formal analysis, methodology, and visualization. H.N. contributed to investigation, methodology, and visualization. O.R. contributed to investigation. G.A. contributed to formal analysis and visualization. C.C., W.S., and M.S. contributed to methodology. P.W. contributed to investigation and methodology. G.D. and D.R. contributed to data curation. K.C., M.R., A.S, S.W.W., T.L., L.M., A.C., J.R. and M.W. contributed resources. Y.A. contributed to methodology, supervision and writing—review and editing. R.T. contributed to conceptualization, funding acquisition, methodology, resources, supervision, and writing— review and editing. All authors: writing-review and editing.

## Prior presentation

This work was previously presented at the Asia-Pacific Vaccine and Immunotherapy Congress (APVIC) 2026 and European Alliance of Associations for Rheumatology Congress (EULAR) 2026.

## Data availability

Single cell RNA sequencing data from the 386.10 study have been deposited in NCBI’s Gene Expression Omnibus (GEO) and are accessible through GEO Series accession number GSE225592 (https://www.ncbi.nlm.nih.gov/geo/query/acc.cgi?acc=GSE225592). Single-cell RNA sequencing data from the AMP RA/SLE study are available via the ARK Portal (10.7303/syn47217489.1) The data are available under controlled access due to data privacy laws. To access the data, users need to complete and submit a signed Data Use Certificate (DUC) to the ARK Portal at https://arkportal.synapse.org/Data%20Access. Flow cytometric and PhenoCycler data supporting the findings of this study may be made available upon reasonable request to the authors, subject to ethics approval and approval by the Australian Arthritis and Autoimmune Biobank Collaborative (A3BC; Access ID: ResID8-PID1).

## Abbreviations

A3BC: Australian Arthritis and Autoimmune Biobank Collaborative
ACPA: Anti-citrullinated peptide autoantibodies
AMP RA/SLE: RA and Systemic Lupus Erythematosus
ANOVA: Analysis of variance
CDAI: Clinical Disease Activity Index
csDMARDs: Conventional synthetic disease modifying anti-rheumatic drugs
DAS28-CRP: Disease activity scores using C-reactive protein
FDR: FDR
HLA: Human leukocyte antigen
MHC: Major histocompatibility complex
NET: Neutrophil extracellular trap
PB: Peripheral blood
PBMCs: Peripheral blood mononuclear cells
RA: Rheumatoid arthritis
RPMI: Roswell Park Memorial Institute medium
SD: Standard deviation
SE: Shared epitope
ST: Synovial tissue
Tem: T effector memory
Tfh: Follicular helper T cells
Th2 Tem: T helper-2 effector memory
Tph: CD4⁺ peripheral helper T cells
UMAP: Uniform manifold approximation and projection

