## Supplementary methods for "Persistent circulating autoreactive PD1⁺TIGIT⁺ peripheral helper T cells reflect synovial lymphoid activity and poor response to conventional disease-modifying anti-rheumatic drugs in early rheumatoid arthritis"

### Hee et al. Supplementary data

#### Methods

#### Spectral flow cytometry

PBMCs cryopreserved in liquid nitrogen were thawed and rested for 2 hours at 37°C in complete RPMI including 10% human AB serum. To prevent non-specific binding, Human TruStain FcX Blocking Reagent (BioLegend) was applied. PE-labelled tetramer was subsequently added at a concentration of 6 µg/mL and incubated for 30 minutes at room temperature, followed by CCR7 BUV805 for an additional 30 minutes, a wash, then a cocktail of surface antibodies incubated for 30 minutes (Supplementary Table 7). LIVE/DEAD FVS700 dye was used to discriminate live from dead cells. All samples were acquired immediately on a spectral flow cytometer (Aurora, Cytek) and analysed using SpectroFlow.

Data analysis included manual (supervised) gating of CD4+ T cells and Cit-vimentin-reactive $CD4^+$ T cells in FlowJo and unsupervised clustering using *Spectre* package in R (13). Boolean gating to identify population overlap employed *CytoML* and *flowWorkspace* in R by constructing an intersection Boolean filter at the T cell parent node within the FlowJo-imported gating set. Overlapping event counts were extracted from the resulting Boolean population. Results were validated in FlowJo. Cit-vimentin-reactive CD4+ T cells were analysed as a single pooled population comprising tetramer-positive cells identified using either tetramer.

#### Unsupervised clustering of flow cytometry data

Before performing FlowSOM clustering, the raw CD4+ data were checked and compensated in FlowJo, then downsampled to 15,000 cells using the Spectre package in R and finally exported as a .FCS file. To prevent the loss of tetramer-positive cells due to their low abundance, these cells were extracted prior to downsampling, exported separately as .FCS files, and subsequently reintegrated into the CD4+ dataset in R.

In R, both $CD4^+$ and tetramer-positive $CD4^+$ T cells .FCS files were loaded, and tetramer-positive cells were reinstated to the CD4+ T cell dataset. The data were then transformed using an arcsinh transformation. To address technical variations between batches, batch controls processed with each batch were used, and batch alignment was performed using the CytoNorm algorithm. Following batch correction, unsupervised clustering was conducted using FlowSOM, and dimensionality reduction was performed with UMAP. Finally, clusters were manually annotated based on marker expression profiles, and clusters containing tetramer-positive $CD4^+$ T cells were separated from $CD4^+$ T cells.

### Single-cell analysis

#### Sample processing

PBMCs were isolated from peripheral blood of participants. Synovial tissues arthroscopic biopsies were collected from participants at the same time as PBMC dissecting into smaller fragments. Both PBC and ST were then cryopreserved in freezing medium (10% DMSO, 50% RPMI-1640, and 40% FCS) until used for assessment. When required, frozen PBMCs were thawed, washed in complete RPMI-1640 media (cRPMI; 1% PSG, 1% NaPyr, 10% FBS) containing 12.5 µg/mL DNase I, and rested in cRPMI containing 6.25 µg/mL DNase I for 30 minutes at 37 °C. Synovial tissues were disaggregated mechanically and enzymatically in digestion buffer (Liberase TL 100 µg/ml and DNase I 100 µg/ml in RPMI) in a waterbath for 30 minutes at 37 °C. Cells from PBMCs and synovial tissues were counting and their viability assessed using Trypan Blue.

PBMC and disaggregated ST cells were labelled with lived dead stains and antibodies for sorting by flow cytometer. Some samples were oligo-tagged (TotalSeq-C0256 anti-human Hashtag 6 or TotalSeq-C0257 anti-human Hashtag 7) and combined prior to sorting. The antibodies used include: Live/Dead Fixable viability stain 700, CD45 APC or FITC, CD3 BUV737, CD4 BUV395, CD8 BV510, CD19 PE, CD56 FITC, CD16 APC/Cy7, CD14 PerCP/Cy5.5, HLA-DR PE/Cy7, CD11c PE-CF594, CD31 AF488, PDPN BV421 and CD90 PE/Cy5. After staining, viable CD45+ leukocytes and fibroblasts (Live CD45- CD3- CD31- PDPN+ cells) were sorted using a FACSAria II cell sorter with a 100 µm nozzle, collecting cells directly into 500 µL of 100% fetal bovine serum (FBS). Samples were run over several days, requiring downstream batch correction

#### Single-cell sequencing data loading and quality control

Data were log-normalized, and highly variable genes were identified for downstream analyses. Scaled expression values were used for principal component analysis. Cell populations were annotated based on canonical lineage-defining gene expression markers. Raw gene expression matrices generated by Cell Ranger (10x Genomics) were imported for each sample individually using Scanpy (v1.11.5). Samples collected from paired peripheral blood mononuclear cells (PBMCs) and synovial tissue (ST) Samples underwent cell hashing demultiplexing prior to analysis. Following demultiplexing, cells from the same patient were merged into a single object per compartment (PBMC and ST).

Quality control (QC) metrics were computed for each cell, including the total UMI count (n_counts), the number of detected genes (n_genes), and the percentage of reads mapping to mitochondrial genes (pct_mito). Cells failing any of the following thresholds were excluded: n_counts < 1,000; n_genes < 500 (PBMC) or n_genes < 200 (ST); pct_mito ≥ 10%. Genes detected in fewer than 3 cells were removed. Doublets were identified and removed using Scrublet as the primary method, with the expected doublet rate scaled proportionally to cell input (0.8% per 1,000 cells, bounded between 2% and 10%). QC outliers exceeding the 99th percentile for both n_genes and n_counts simultaneously were flagged as a supplementary doublet criterion. Cells called as doublets by Scrublet were excluded prior to downstream analysis.

**Single- cell data integration**

Cleaned, per-sample AnnData objects were concatenated across all patients and compartments (PBMC and ST). Raw count matrices were retained as the input for integration, as scVI does not require prior log-transformation or PCA.

Batch correction and data integration were performed using scVI (scvi-tools) (32). The model was configured with batch_key="batch" to correct for sequencing batch effects across three library preparation batches, with patient identity additionally included as a categorical covariate to account for inter-individual variation. The scVI model was trained with a negative binomial gene likelihood, 2 encoder/decoder layers, a latent space dimensionality of 50, and a maximum of 200 training epochs. The latent representation was stored and used for all downstream analyses.

**Cell clustering and pan-T cell annotation**

A k-nearest neighbour graph was constructed from the scVI latent embedding using n_neighbors=30. Uniform Manifold Approximation and Projection (UMAP) was computed on this graph for two-dimensional visualisation. Leiden community detection was applied at resolution=0.5 to define unsupervised cell clusters.

Raw counts were normalised to 10,000 counts per cell and log1p-transformed for marker gene scoring. Gene module scores were computed for each of the following cell populations using sc.tl.score_genes: Pan T cells (CD3D, CD3E, CD247, TRAC, TRBC1, TRBC2, LCK), B cells (MS4A1, CD79A, CD79B, CD74, HLA-DRA, CD37), plasma cells (MZB1, XBP1, SDC1, JCHAIN, IGHG1, IGKC), classical monocytes (LYZ, S100A8, S100A9, FCN1, CTSS), non-classical monocytes (FCGR3A, LST1, MS4A7, IFITM3, CTSS), macrophages (C1QA, C1QB, C1QC, APOE, LGALS3, CTSS), cDC2 (FCER1A, CD1C, CLEC10A, ITGAX, HLA-DRA, CD74), cDC1 (CLEC9A, BATF3, IRF8, XCR1, ITGAX), plasmacytoid dendritic cells (IRF7, TCF4, IL3RA, SERPINF1) and fibroblasts (COL1A1, COL1A2, DCN, LUM, COL3A1, PRG4). Mean scores per Leiden cluster were computed and labels assigned using a hierarchical decision rule prioritising: (1) T cells (pan-T score > 0.05), (2) plasma cells (plasma score > B cell score + 0.03), (3) B cells (B score > 0.05), (4) macrophages, (5) monocytes, (6) dendritic cell subtypes, and (7) stromal populations. Clusters not meeting any threshold were labelled as Unknown. Cells pan-T were subsetted from the integrated object to generate a pan-T cell AnnData object for downstream T cell subtype analysis.

To improve transcriptomic discrimination between CD4 and CD8 T cells, we implemented an overlap-aware classification framework using ADT-defined reference populations. ADT-labelled CD4 and CD8 T cells were first classified based on their transcriptomic profiles, revealing a subset of cells with discordant protein and RNA identities. Specifically, some ADT-defined CD8 T cells exhibited transcriptional profiles more like canonical CD4 T cells ("hard CD8"), while a reciprocal population of ADT-defined CD4 T cells displayed CD8-like transcriptional features ("hard CD4"). These discordant populations were analysed separately from the canonical CD4 and CD8 populations to identify genes associated with transcriptional overlap between CD4 and CD8 states. Genes derived from both canonical and discordant comparisons were combined into an expanded feature set and used to train a LASSO-regularised logistic regression classifier using the glmnet package. ADT-defined CD4 and CD8 T cells from Batch 2 were randomly partitioned into training (70%) and test (30%) sets at the cell level in a class-stratified manner. LASSO regularisation ($\alpha = 1$) was used to reduce overfitting and perform embedded feature selection by shrinking uninformative coefficients towards zero. The regularisation parameter ($\lambda$) was selected by five-fold cross-validation within the training set, and the model corresponding to the minimum cross-validation error (lambda.min) was selected. The resulting model generated a continuous CD4–CD8 transcriptional score and corresponding class probabilities for all T cells. To define high-confidence classifications, a range of probability thresholds were evaluated to maximise classification coverage while maintaining a minimum accuracy of 97%. Cells with probabilities above the upper threshold were assigned as CD4, cells below the complementary lower threshold were assigned as CD8, and intermediate cells were classified as ambiguous. Model performance was then evaluated on held-out test cells. The fitted classifier was subsequently applied to all T cells across the full dataset, enabling annotation of CD4 and CD8 identity in batches lacking ADT measurements.

**Spatial proteomics analysis**

Marker intensity data were compiled into an AnnData object, where rows represented cells and columns represented marker fluorescence intensities. Cells were annotated with metadata including tissue region, slide, patient ID, paired baseline and 6-month sample IDs. The lowest 1% of cells by overall fluorescence intensity and cell size were omitted. Z-score normalisation was applied to minimise slide batch effects. Additional filtering removed the top 1% of cells with the highest summed fluorescence intensity and the top 1% of cells positive for the greatest number of markers. Leiden unsupervised clustering was performed and used as the basis for downstream cell annotation (14).

### Supplementary Tables

Supplementary Table 1. Population-specific offset using parent cell count for citrullinated vimentin-specific CD4+ T cells clustered using supervised gating.

| | Offset |
|---|---|
| Tet+ CD27+ | Total Tet+ |
| Tet+ CXCR3 CCR4 DP | Tet+ CD4 Total memory |
| Tet+ CXCR3+CCR4- | Tet+ CD4 Total memory |
| Tet+ Th1 | Tet+ CD4 Total memory |
| Tet+ Th1 Th17 | Tet+ CD4 Total memory |
| Tet+ CXCR3-CCR4+ | Tet+ CD4 Total memory |
| Tet+ Th2 | Tet+ CD4 Total memory |
| Tet+ Th17 | Tet+ CD4 Total memory |
| Tet+ GPR56+ | Total Tet+ |
| Tet+ KLRG1+ | Total Tet+ |
| Tet+ non-Treg | Total Tet+ |
| Tet+ Tfh | Tet+ non-Treg |
| Tet+ Tph | Tet+ non-Treg |
| Tet+ PD1 TIGIT DP | Total Tet+ |
| Tet+ PD1+ | Total Tet+ |
| Tet+ Tcm | Total Tet+ |
| Tet+ Tem | Total Tet+ |
| Tet+ Temra | Total Tet+ |
| Tet+ TIGIT+ | Total Tet+ |
| Tet+ Tn | Total Tet+ |
| Tet+ Treg | Total Tet+ |
| Tet+ Tfr | Tet+ Treg |
| Tet+ CD4 Total memory | Total Tet+ |
| Tet+ CD39+ | Total Tet+ |
| Tet+ Treg CD39+ | Tet+ Treg |

Supplementary Table 2. Population-specific offset using parent cell count for CD4+ T cells clustered using supervised gating.

| | Offset |
|---|---|
| CD4+ CD27+ | Total CD4+ |
| CD4+ CXCR3 CCR4 DP | CD4+ CD4 Total memory |
| CD4+ CXCR3+CCR4- | CD4+ CD4 Total memory |
| CD4+ Th1 | CD4+ CD4 Total memory |
| CD4+ Th1 Th17 | CD4+ CD4 Total memory |
| CD4+ CXCR3-CCR4+ | CD4+ CD4 Total memory |
| CD4+ Th2 | CD4+ CD4 Total memory |
| CD4+ Th17 | CD4+ CD4 Total memory |
| CD4+ GPR56+ | Total CD4+ |
| CD4+ KLRG1+ | Total CD4+ |
| CD4+ non-Treg | Total CD4+ |
| CD4+ Tfh | CD4+ non-Treg |
| CD4+ Tph | CD4+ non-Treg |
| CD4+ PD1 TIGIT DP | Total CD4+ |
| CD4+ PD1+ | Total CD4+ |
| CD4+ Tcm | Total CD4+ |
| CD4+ Tem | Total CD4+ |
| CD4+ Temra | Total CD4+ |
| CD4+ TIGIT+ | Total CD4+ |
| CD4+ Tn | Total CD4+ |
| CD4+ Treg | Total CD4+ |
| CD4+ Tfr | CD4+ Treg |
| CD4+ CD4 Total memory | Total CD4+ |
| CD4+ CD39+ | Total CD4+ |
| CD4+ Treg CD39+ | CD4+ Treg |

Supplementary Table 3. Population-specific offset using parent cell count for citrullinated vimentin-specific CD4+ T cells clustered using unsupervised gating.

| | Offset |
|---|---|
| Tet+ CXCR3+ Tph | Total Tet+ |
| Tet+ GPR56+ Temra | Total Tet+ |
| Tet+ GPR56+ Tph | Total Tet+ |
| Tet+ KLRG1+ anergic | Total Tet+ |
| Tet+ Naive | Total Tet+ |
| Tet+ Naive Tregs | Tet+ Naive |
| Tet+ Tem | Total Tet+ |
| Tet+ Tfh | Total Tet+ |
| Tet+ Th1 Tem | Total Tet+ |
| Tet+ Th17.2 | Total Tet+ |
| Tet+ Th2 | Total Tet+ |
| Tet+ Tph | Total Tet+ |
| Tet+ Treg | Total Tet+ |

Supplementary Table 4. Population-specific offset using parent cell count for CD4+ T cells clustered using unsupervised gating.

| | Offset |
|---|---|
| CD4+ CXCR3+ Tph | Total CD4+ |
| CD4+ GPR56+ Temra | Total CD4+ |
| CD4+ GPR56+ Tfh | Total CD4+ |
| CD4+ GPR56+ Tph | Total CD4+ |
| CD4+ KLRG1+ anergic | Total CD4+ |
| CD4+ Naive | Total CD4+ |
| CD4+ Naive Tregs | CD4+ Naive |
| CD4+ Tem | Total CD4+ |
| CD4+ Tfh | Total CD4+ |
| CD4+ Th1 Tem | Total CD4+ |
| CD4+ Th17.2 | Total CD4+ |
| CD4+ Th2 | Total CD4+ |
| CD4+ Tph | Total CD4+ |
| CD4+ Treg | Total CD4+ |

Supplementary Table 5. Characteristics of cohort 2 donors.

| | 396.10 study, N = 22 | AMP-RA-SLE, N = 4 |
|---|---|---|
| Age at baseline, mean (SD) | 54.5 (17.1) | 67.8 (3.9) |
| Symptom duration at baseline (weeks), median (IQR) | 14.3 (9.3 – 25.5) | 35.5 (15.8 – 59.5) |
| Sex, n (%) | | |
| Female | 15 (68.2) | 4 (100) |
| Male | 7 (31.8) | 0 (0) |
| HLA-DR-SE+, n (%) | 15 (68.2%) | 3 (75%, 1 missing) |
| ACPA, n (%) | | |
| Present | 15 (68.2) | 3 (75.0) |
| Absent | 7 (31.8) | 1 (25.0) |
| Pathotype, n (%) | | |
| Diffuse-myeloid | 9 (40.9) | 0 (0) |
| Lympho-myeloid | 9 (40.9) | 4 (100.0) |
| Pauci-immune | 4 (18.2) | 0 (0) |
| DAS-CRP at baseline, median (IQR) | 5.3 (4.6 – 5.1) | 5.0 (4.1 – 5.8) |
| DAS-CRP at 6 months, median (IQR) | 1.8 (1.5 – 2.4) | - |

ACPA = Anti-citrullinated protein antibodies, DAS-CRP = disease activity score using C-reactive protein, DMARDS = disease modifying anti-rheumatic drugs, , HLA-DR-SE+ = HLA-DRB1 shared epitope positive, IQR = interquartile range, RF = rheumatoid factor, SD = standard deviation. * DAS-CRP estimated from CDAI.

Supplementary Table 6. List of differentially expressed genes between CD4$^+$ TIGIT$^+$ Tph and in CD4$^+$ TIGIT$^-$ Tph/Tfh cells. Genes with log2 fold change > 0.25, expressed in at least 25% of cells, and adjusted p-value ≤ 0.05 were considered significant. Excel file uploaded.

Supplementary Table 7. Antibody cocktail used in flow panel

| Marker | Fluorochrome |
|---|---|
| Live/Dead | FVS700 |
| Tetramer | PE |
| Tetramer | APC |
| CD3 | BUV737 |
| CD4 | BUV395 |
| CD8 | BUV496 |
| CD25 | PE-Cy7 |
| CD127 | BV421 |
| PD1 | BB700 |
| KLRG1 | PE-Dazzle594 |
| CCR6 | BV480 |
| CXCR3 | PE-Cy5 |
| CCR4 | BV605 |
| GPR56 | APC-Vio770 |
| CCR7 | BUV805 |
| CD45RA | FITC |
| CD27 | BV650 |
| TIGIT | BV786 |
| CXCR5 | BV711 |
| CD39 | NovaFluor Blue610-70S |

### Supplementary Figures

Supplementary Figure 1. Flow cytometry gating strategy for identification of CD4$^+$ T cell and citrullinated-vimentin reactive CD4$^+$ T cell subsets. A: total CD4+ T cells; B: tetramer gated CD4+ T cells

A

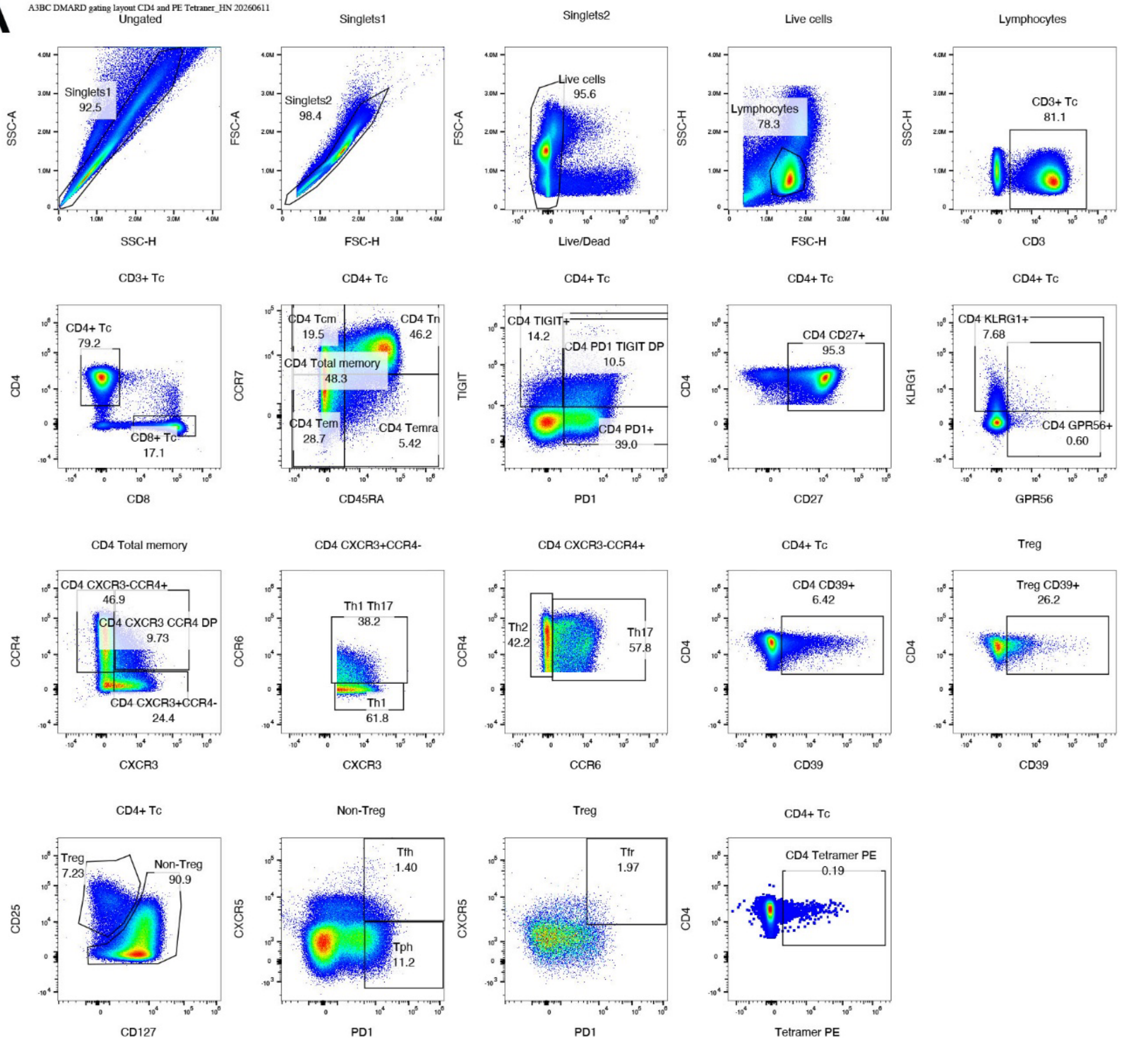

**B**

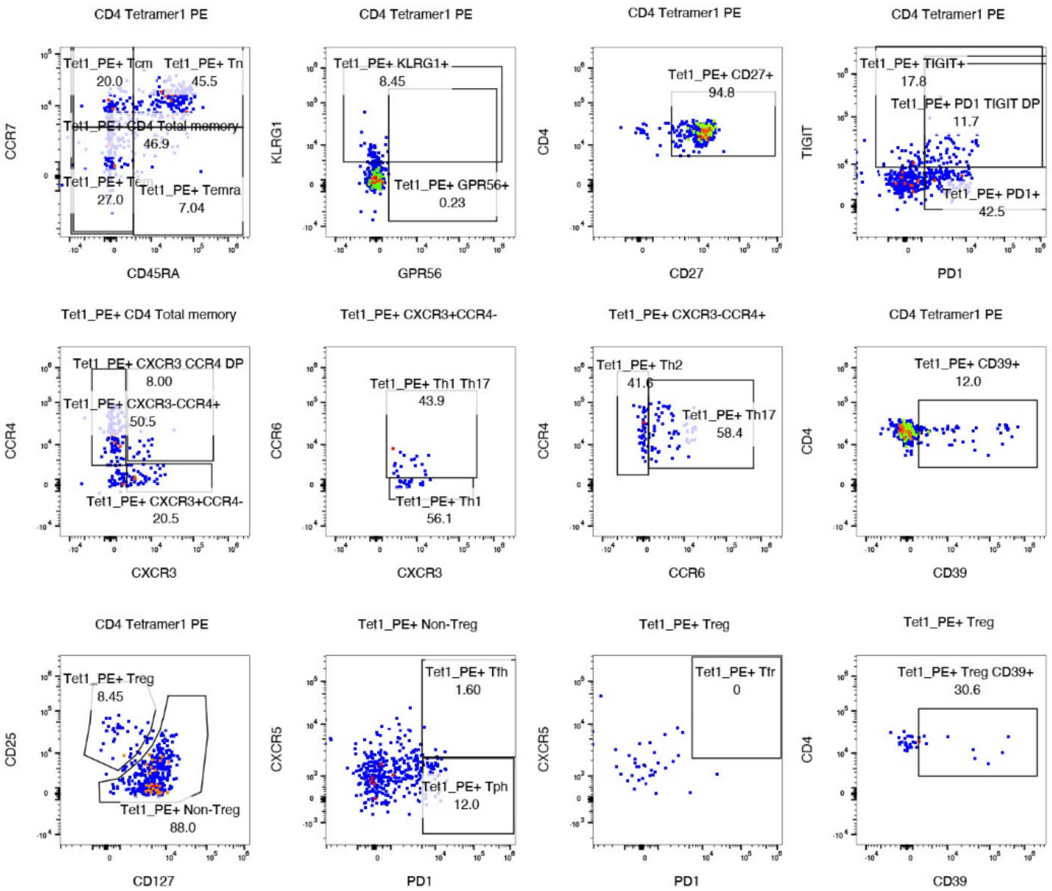


Supplementary Figure 2. Dot plot of protein marker expression for follicular B cells, plasma cells, Treg and Tfh/Tph in PhenoCycler images.

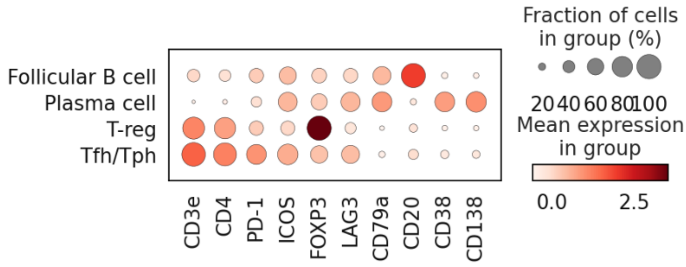

Supplementary Figure 3. Flow-cytometry plots demonstrating gating strategy to obtain frequencies of CD4$^{+}$ Tph TIGIT$^{+}$ and CD4$^{+}$ Tph TIGIT$^{-}$ from non-regulatory CD4$^{+}$ T cells, and non-Tph CD4$^{+}$ PD1$^{+}$ TIGIT$^{+}$ DP from total CD4$^{+}$ cells.

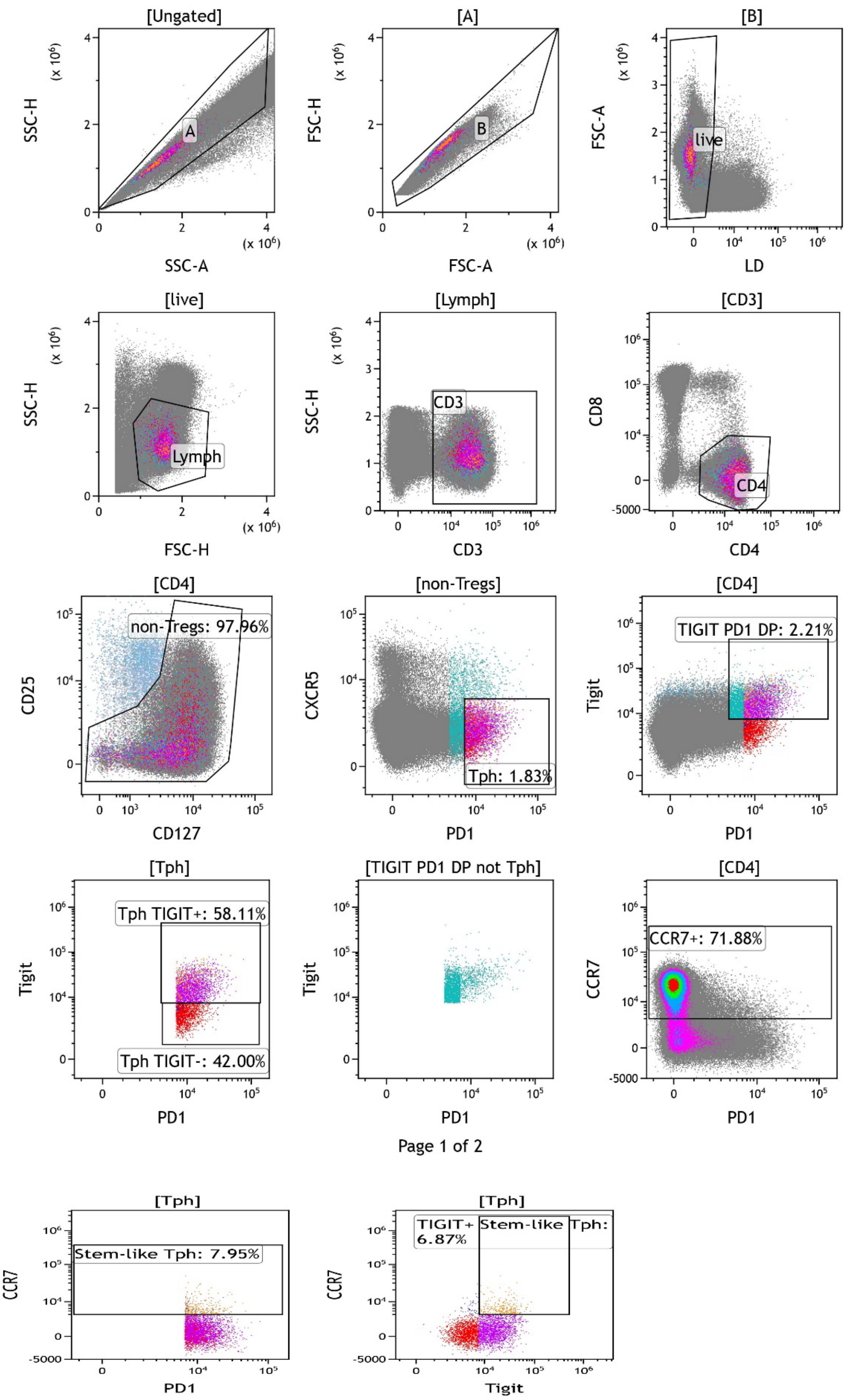

Supplementary Figure 4. (A-B) Bar plots demonstrating the frequency of CD4+ T cells clustered using unsupervised clustering stratified by disease severity. Grey shaded areas represent the 95% confidence interval. Low = Low disease activity at point of observation, Mod/High = Moderate or high disease activity at point of observation, Remission = Remission at point of observation. (C) Spline trajectories of differentially expressed unsupervised clustered CD4$^{+}$ Tfh cells over time (baseline, 6 months and 12 months) stratified by stable remission and non-remission.

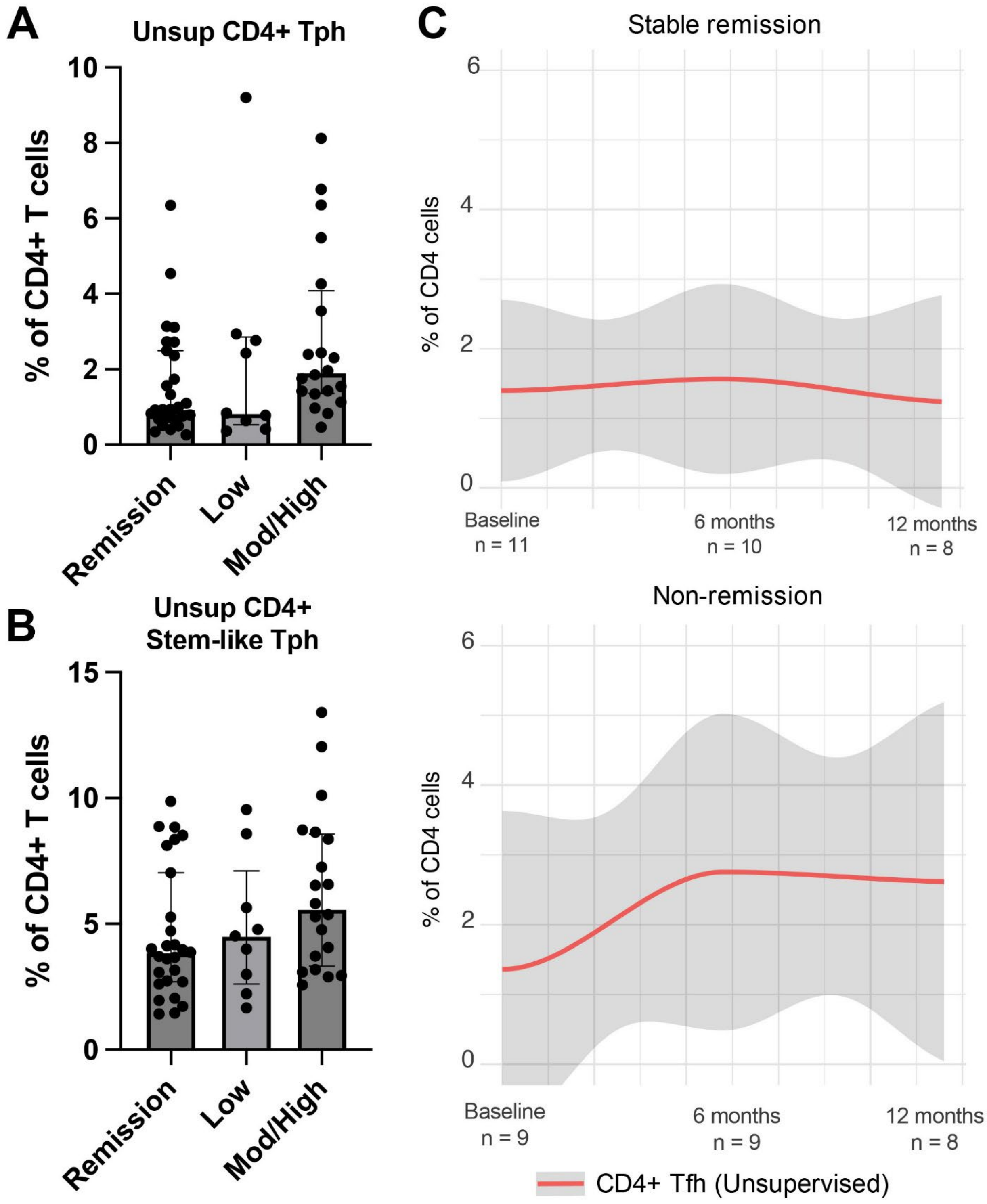

Supplementary Figure 5. UMAP visualisation of functional state markers expressed in CD4+ T cells.

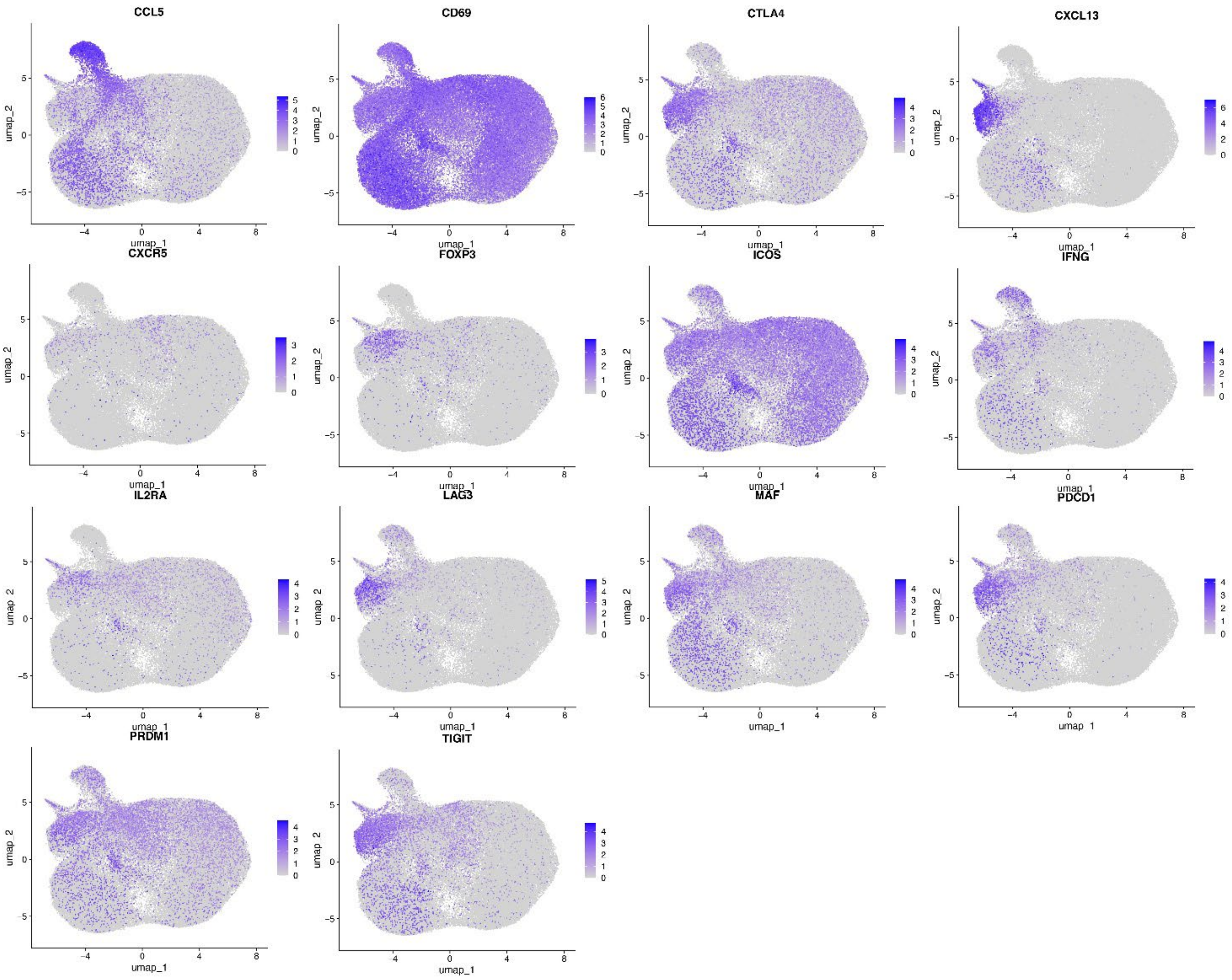

Supplementary Figure 6. Violin plots showing the distribution of the number of detected genes (nFeature_RNA), total RNA counts (nCount_RNA), percentage mitochondrial transcripts (percent.mt), and percentage ribosomal transcripts (percent.ribo) in TIGIT+ and TIGIT− Tph cells.

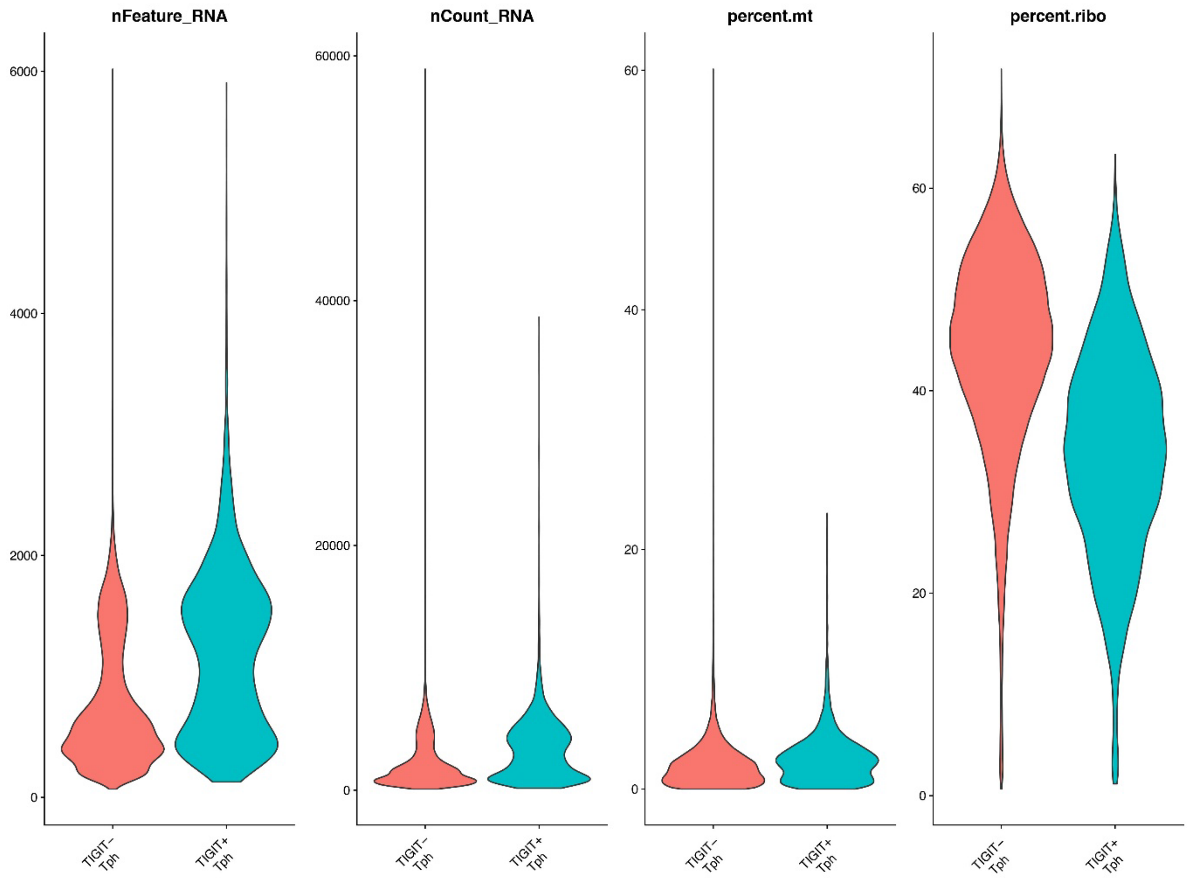

Supplementary Figure 7. PPI network (confidence score of 0.90 and no interactors allowed) linking proteins encoded by upregulated genes in CD4$^{+}$ TIGIT$^{+}$ Tph (in grey) and in CD4$^{+}$ TIGIT$^{-}$ Tph (in blue). Nodes represent proteins, and edges represent known protein-protein interactions. The size of each node indicates the degree of connectivity of the corresponding protein within the network, with larger nodes representing hub proteins.

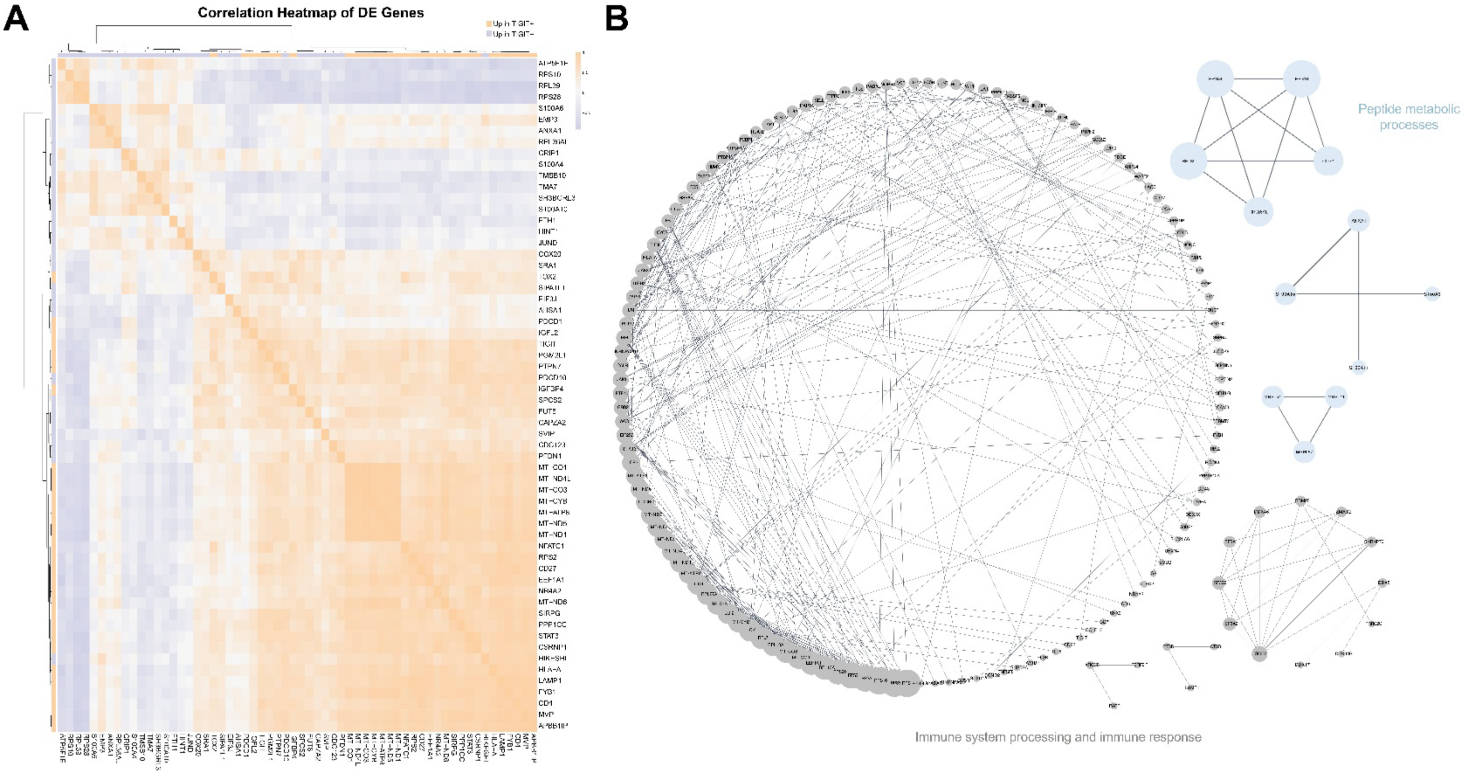
